# Barcode-free prediction of cell lineages from scRNA-seq datasets

**DOI:** 10.1101/2022.09.20.508646

**Authors:** A.S. Eisele, M. Tarbier, A.A. Dormann, V. Pelechano, D.M. Suter

**Affiliations:** Ecole Polytechnique Federale de Lausanne, School of Life Sciences, Institute of Bioengineering; Science for Life Laboratory, Department of Microbiology, Tumor and Cell Biology, Karolinska Institute, Solna, Sweden

## Abstract

The integration of lineage tracing with scRNA-seq has transformed our understanding of gene expression heritability during development, regeneration, and disease. However, lineage tracing is technically demanding and most existing scRNA-seq datasets are devoid of lineage information. Here we introduce <u>G</u>ene <u>E</u>xpression <u>M</u>emory-based <u>L</u>ineage Inference (GEMLI), a computational pipeline allowing to predict cell lineages over several cell divisions solely from scRNA-seq datasets. GEMLI leverages genes displaying conserved expression levels over cell divisions, and allows i.a. identifying cell lineages in a broad range of cultured cell types, in intestinal organoids, and in crypts from adult mice. GEMLI recovers GO-terms enriched for heritable gene expression, allows to discriminate symmetric and asymmetric cell fate decisions and to reconstruct individual cellular structures from pooled scRNA-seq datasets. GEMLI considerably extends the pool of datasets from which lineage information can be obtained, thereby facilitating the study of gene expression heritability in a broad range of contexts. GEMLI is available at (https://github.com/UPSUTER/GEMLI).

## Introduction

The quantification of mRNA levels in single cells has revealed extensive transcriptome heterogeneity not only between but also within a given cell type ^1–6^. Heterogeneous mRNA levels for given genes are often transient, but the expression levels of some genes can be preserved through cell division ^7^. The existence of such gene expression memory is of major fundamental and biomedical interest, as it is involved in cancer resistance to drugs ^8–14^, disease onset ^11,15^, and differentiation trajectories in development, homeostasis and regeneration ^16–21^. To study gene expression over cell divisions, scRNA-seq has been combined with different lineage tracing techniques. One is cellular barcoding, which involves the introduction of heritable, expressed DNA barcodes in individual cells, which can be retrieved in scRNA-seq data ^17,18,22–26^. Other approaches involve time-lapse microscopy in combination with smFISH, and the use of dedicated microfluidic devices or sister-cell picking before scRNA-seq ^27–31^. Bulk sequencing approaches can also be used by comparing the variability of gene expression of individually cultured clones grown for several days to their population of origin ^8,10,15^.

All of these techniques are powerful but present important technical limitations. Microfluidics, time-lapse imaging, sister cell picking, and bulk sequencing based approaches depend on dedicated microfluidic devices, extensive imaging and cell tracking, and/or extensive cell culture, respectively. Likewise, lentiviral cellular barcoding can be limited by cells that are hard to transduce, requires genetic engineering and a cellular barcoding library of adequate size whose barcodes are strongly expressed in the cells of interest. Approaches for in vivo cellular barcoding involve complex procedures such as tamoxifen-induced Cre recombination or CRISPR/Cas9-induced scarring, which can result in the asynchronous generation of barcodes among founder cells or variable barcode quality, respectively. This makes these approaches not easily amenable to studying small cell lineages. Finally, the analysis of cellular barcoding data is cumbersome and the quality of cell lineage assignment is variable.

Here, we introduce <u>G</u>ene <u>E</u>xpression <u>M</u>emory-based <u>L</u>ineage Inference or GEMLI, a computational tool that allows identifying cell lineages (i.e. cells sharing a common ancestor) based solely on scRNA-seq data. We analyze new and publicly-available lineage-annotated scRNA-seq datasets of multiple cell types and identify genes that are particularly stable in their expression levels across several cell divisions (memory genes). We determine common characteristics of memory genes to enrich such genes in scRNA-seq data. This allows us to predict cell lineages de novo using an iterative hierarchical clustering approach, in a broad range of cell types cultured in vitro, in intestinal organoids and in intestinal crypts from mice. We show that de novo lineage prediction can identify cell type-specific heritable gene expression programs, symmetric vs asymmetric cell fate decisions, gene expression programs specific to cells switching to a particular fate, and correctly assign cells to individual intestinal organoids or crypts in merged scRNA-seq datasets.

## Results

### Cells belonging to the same lineage tree are similar in their gene expression profiles

To assess the stability of gene expression over cell division, we generated a lineage-annotated scRNA-seq dataset of mouse embryonic stem cells (mESCs). We used the LARRY barcoding library ^17^ to transduce mESCs at low multiplicity of infection, sorted barcoded cells (Extended Data Fig.1a-c) and grew them for 48h before scRNA-seq. In addition, we used public lineage-annotated datasets of other cell types: primary mouse embryonic fibroblasts (MEF) ^18^, primary mouse CD8+ T-lymphocytes (CD8), lymphocytic leukemia cells (L1210) (both ^29^), primary mouse hematopoietic stem and progenitor cells (HSPC, more precisely LK and LSK subsets) ^17^, hematopoietic stem cells (HSC) ^27^, melanoma cells (WM989) ^9^, as well as intestinal crypts and organoids ^32^. These datasets were generated using different lineage assignment and scRNA-seq platforms and encompassed cells in self-renewal or differentiation conditions grown for 2-14 days (Table 1).

**Table 1:** ScRNA-seq datasets with annotated cell lineage information.

| Cell type | Source | State | Lineage info | Culture time | Divisions | Sequencing |
| --- | --- | --- | --- | --- | --- | --- |
| mESC | Own data | Self-renewal | LARRY | 48h | Avg. 2-3 | 10X |
| MEF | Biddy et al. | Steady-state | CellTag V1 | 48h | Unknown | 10X |
| MEF | Biddy et al. | Reprogramming | CellTag V1-V3 | 48h-28 days | Unknown | 10X |
| CD8 | Kimmerling et al. | Activated | Microfluidic | 4-cell | 2 | SmartSeq |
| L1210 | Kimmerling et al. | Steady-state | Microfluidic | 4-cell | 2 | SmartSeq |
| HSPC | Weinreb et al. | Differentiation | LARRY | 48h- 6 days | Avg. 3 | InDrop |
| HSC | Wehling et al. | Differentiation | Cell picking | 2-cell | 1 | Adapted SCRB-seq |
| WM989 | Harmange et al. | Steady-State | Barcoding | 12-14 days | Avg. 5 | 10X |
| Intestinal organoid | Bues et al. | Differentiation | DisCo | 3-6 days | Unknown | Custom |
| Intestinal crypt | Bues et al. | Steady-State | DisCo | None | Unknown | Custom |

To quantify heritability of gene expression, we analyzed data from five cell types (Supplementary Table 1) grown over 48h or 2-3 divisions. We observed a higher correlation of the whole transcriptome for cells from the same lineage as compared to randomly sampled cells for all datasets (Fig.1a-b). Consistent with previous findings that sister cells often exhibit similarities in their transcriptional activity ^7^, RNA velocity momenti ^33^ of related cells were more correlated than in random cells (Fig.1c and Extended Data Fig.2a). Finally, related cells expressed a similar number of genes (complexity; Fig.1d and Extended Data Fig.2b) and were often in the same cell cycle phase (Fig.1e and Extended Data Fig.2c). These findings are consistent with the existence of a gene expression memory that spans several cell divisions in both self-renewing and differentiating cells.

**Figure 1:**
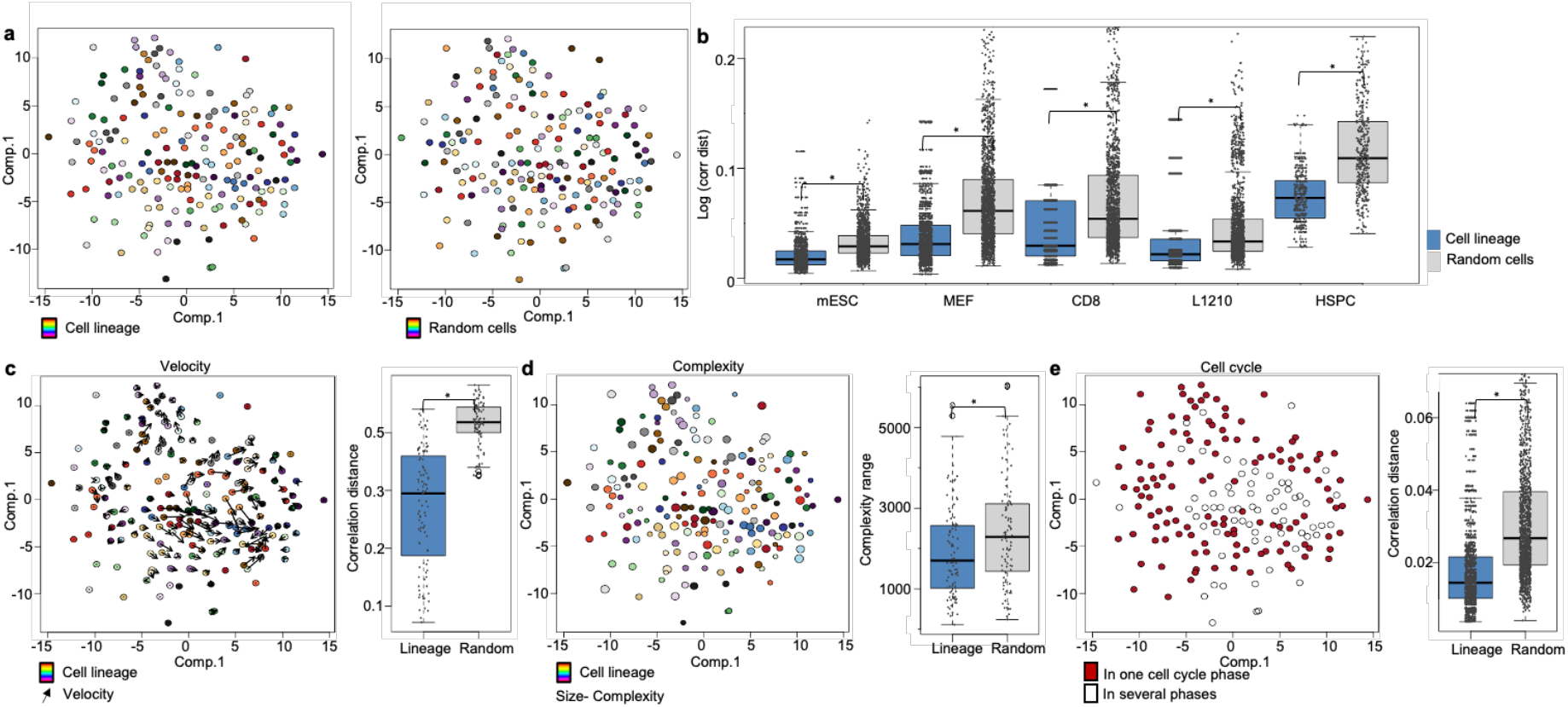
Gene expression memory at the whole transcriptome level. a) tSNE embedding based on a PCA of exonic and intronic reads of the lineage-annotated (left; colors) or randomly-annotated (right) mESC scRNA-seq dataset. Selection on cell lineages of 3 members for ease of visualization. b) Comparison of log (correlation distance) in the different analyzed cell types for cells belonging to the same cell lineage (blue), and randomly sampled cells (gray) in the same dataset (100 repetitions). Boxes: intervals between the 25th and 75th percentile and median (horizontal line). Error bars: 1.5-fold the interquartile range or the closest data point when no data point is outside this range. c-e) tSNE embedding on a PCA of lineage-annotated mESCs as in a) including velocity vectors (c), size as complexity quantile (complexity = number of genes expressed) (d), or colored according to the completeness of a lineage in one cyclone-assigned cell-cycle phase (e). Boxplots are the respective quantification of similarity in related and randomly sampled cells (as correlation distance or complexity range) for cell lineages of 3-5 members. Boxplots as in b. Statistical significance for (b-e) was tested using a Mann-Whitney U-test (p= 0.05).

### Thousands of genes display stable gene expression levels through cell division

We next wanted to identify genes based on their stability in gene expression across cell division, i.e. with or without memory. We reasoned that genes whose expression changes little over cell divisions will show a low variability in mean expression within each given lineage, while the variability in mean expression across the different lineages will be high (Fig.2a). To find such memory genes, we compared the gene-wise variability (coefficient of variation squared (CV^2^)) of the mean level of expression across cell lineages to the distribution in repeated (20x) random samples of matched sizes in the same population to calculate a p-value. We defined memory genes as genes with significantly higher (p<= 0.05) variability in mean expression across cell lineages than across random samples. We identified memory genes in all datasets (on average 3320 +/-S.D. 896, Fig.2b). Using different criteria to define memory genes (see Methods) resulted in strongly overlapping genesets (Extended Data Fig.3a-b). Memory genes were partly shared across cell types, with 968 genes found in >3 cell types, and 85 genes found in all cell types (Fig.2c). Other memory genes (1270+/-509) were cell type-specific (Fig.2c). This suggests that a subset of memory genes drives overall transcriptomic similarity of cell lineages.

**Figure 2:**
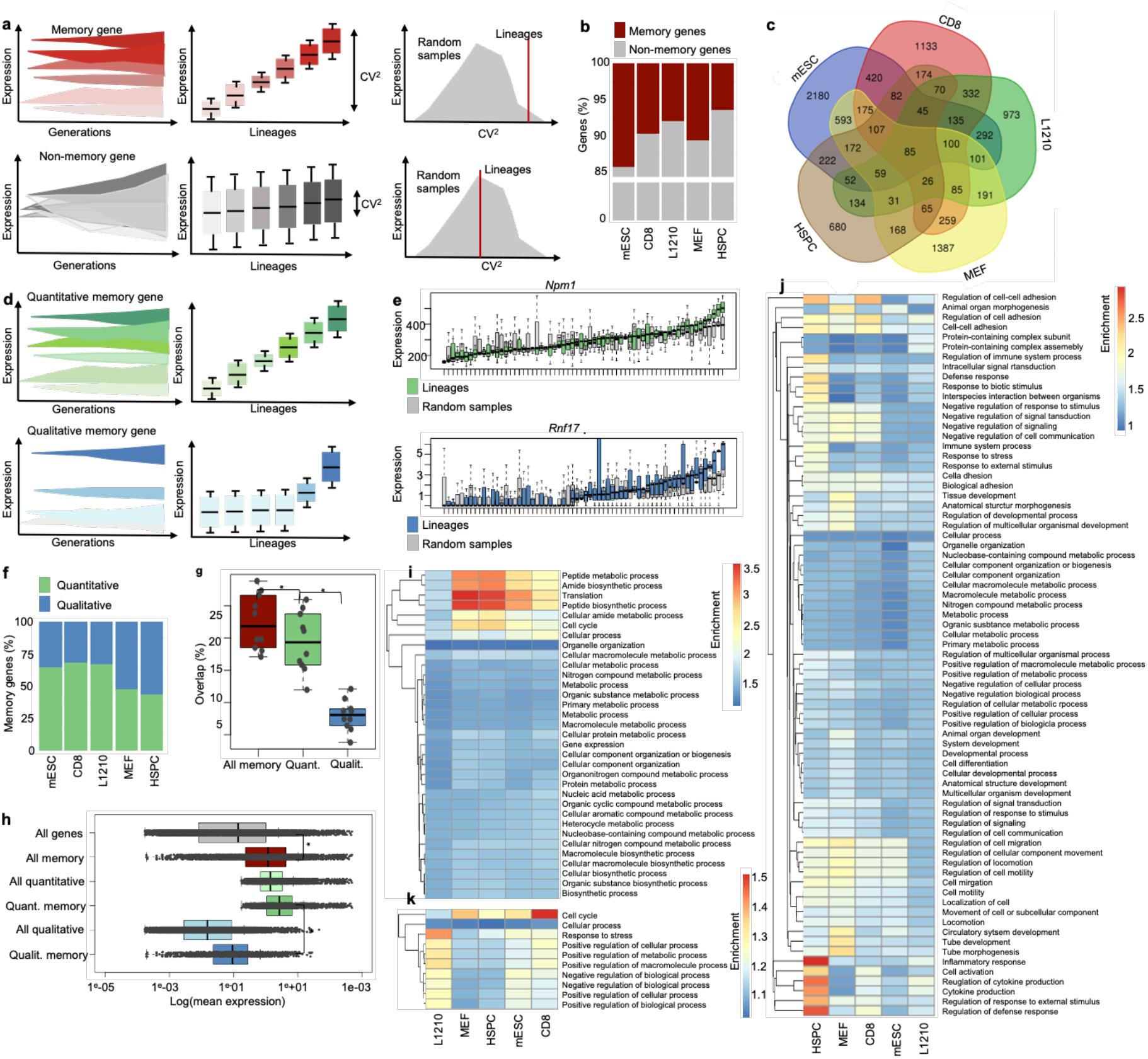
Memory genes with similar characteristics across cell types drive transcriptomic similarity within cell lineages. a) Illustration of memory gene and non-memory gene expression across cell lineages. b) Percentage of genes categorized as memory genes in each cell type. c) Venn diagrams showing the overlap of all memory genes in the different cell types. d) Illustration of quantitative and qualitative memory gene expression across cell lineages. e) Expression level distributions of a quantitative memory gene (*Npm1*), and a qualitative memory gene (*Fabp1*). Blue: cell lineages; Gray: random samples. Boxes: intervals between the 25th and 75th percentile and median (horizontal line). Error bars: 1.5-fold the interquartile range or the closest data point when no data point is outside this range. f) Percentage of quantitative and qualitative memory genes in different cell types. g) Pairwise overlap of memory genes between different cell types. Boxplot as in e. h) Expression levels of the different gene categories. Boxplot as in e. i-j) Heatmap of combined top 20 GO-term enrichment in quantitative (i) and qualitative (j) memory genes of the different cell types. Note that a high overlap of top20 GO-terms in quantitative memory genes results in fewer represented GO terms. k) Heatmap of the highest shared top 100 GO-terms of high memory genes in the different cell types. Significance for g and h was tested using a Mann-Whitney U-test (p= 0.05).

### Memory genes share characteristic expression distributions across cell types

We next classified memory genes with respect to their expression distributions. We defined genes that are expressed in most cells of a given cell population as quantitative genes and genes that are generally lowly expressed but abundant in few cells as qualitative genes (skewness <=1, and >1, respectively). We found memory genes in both categories (Fig.2d-f). Qualitative memory genes were highly expressed only in a small number of cell lineages. Quantitative memory genes were widely expressed, but showed very conserved gene expression levels within individual cell lineages. The fraction of both gene categories was similar across datasets (Fig.2f). Quantitative memory genes were highly shared across cell types, highly-expressed and related to housekeeping functions, while most qualitative memory genes were cell type-specific (Fig.2g-j, Supplementary table 2-4). Cell cycle-related GO terms were among the top enriched terms in shared memory genes (Fig.2k), in line with lineages being often in the same cell cycle phase (Fig.1e).

**Table 2:** Filtering thresholds for the MEF datasets

|  | Mitochondrial % | reads |
| --- | --- | --- |
| D0_2 | 1-14 | 2000 |
| D0 | 1,2-7,5 | 7000 |
| D3_2 | 3-14 | 6000 |
| D3 | 2-10 | 6000 |
| D6_2 | 3-14 | 6000 |
| D6 | 3-20 | 4000 |
| D9_2 | 2-18 | 6000 |
| D9 | 2-18 | 6000 |
| D12_2 | 2-18 | 4000 |
| D12 | 2-18 | 3000 |
| D15_2 | 2-14 | 2000 |
| D15 | 2-18 | 4000 |
| D21_2 | 2-20 | 2000 |
| D21 | 2-18 | 4000 |
| D28_2 | 3-18 | 2500 |
| D28 | 3-18 | 3000 |

### Memory genes allow to predict cell lineages across related cell types

We next wondered whether memory genes could be used to predict lineages within the same dataset. A simple hierarchical clustering based on memory genes grouped only a fraction of related cells (Extended Data Fig.4a). We therefore developed a repetitive, iterative hierarchical clustering approach to predict cell lineages using a given cell type-specific set of memory genes (Fig.3a). Briefly, cells are clustered iteratively on random subsets of the memory genes until being assigned to a cluster of 2-3 cells. By repeating this clustering, every cell pair can be assigned a level of confidence for belonging to the same lineage based on the number of times it has clustered together across individual predictions. When applying this repetitive iterative clustering to different datasets, the precision (true positive (TP)/(TP+ false positive (FP))) and sensitivity (TP/(TP+ false negative (FN))) of the predictions reached an average of 84%+/-12%, and 39%+/-15%, respectively, at a level of confidence of 30 (Fig.3b-f, see Extended Data Fig.4B-E for other confidence levels and datasets). Repetitive iterative clustering using all genes also improved predictions as compared to a simple clustering, but was less powerful than using memory genes (Fig.3b, d-f Extended Data Fig.4a, c-e). Memory genes called using other methods, including a mutual information maximizer or ANOVA F-test approach, did not further improve predictions (see Methods, Extended Data Fig.4f-g).

**Figure 3:**
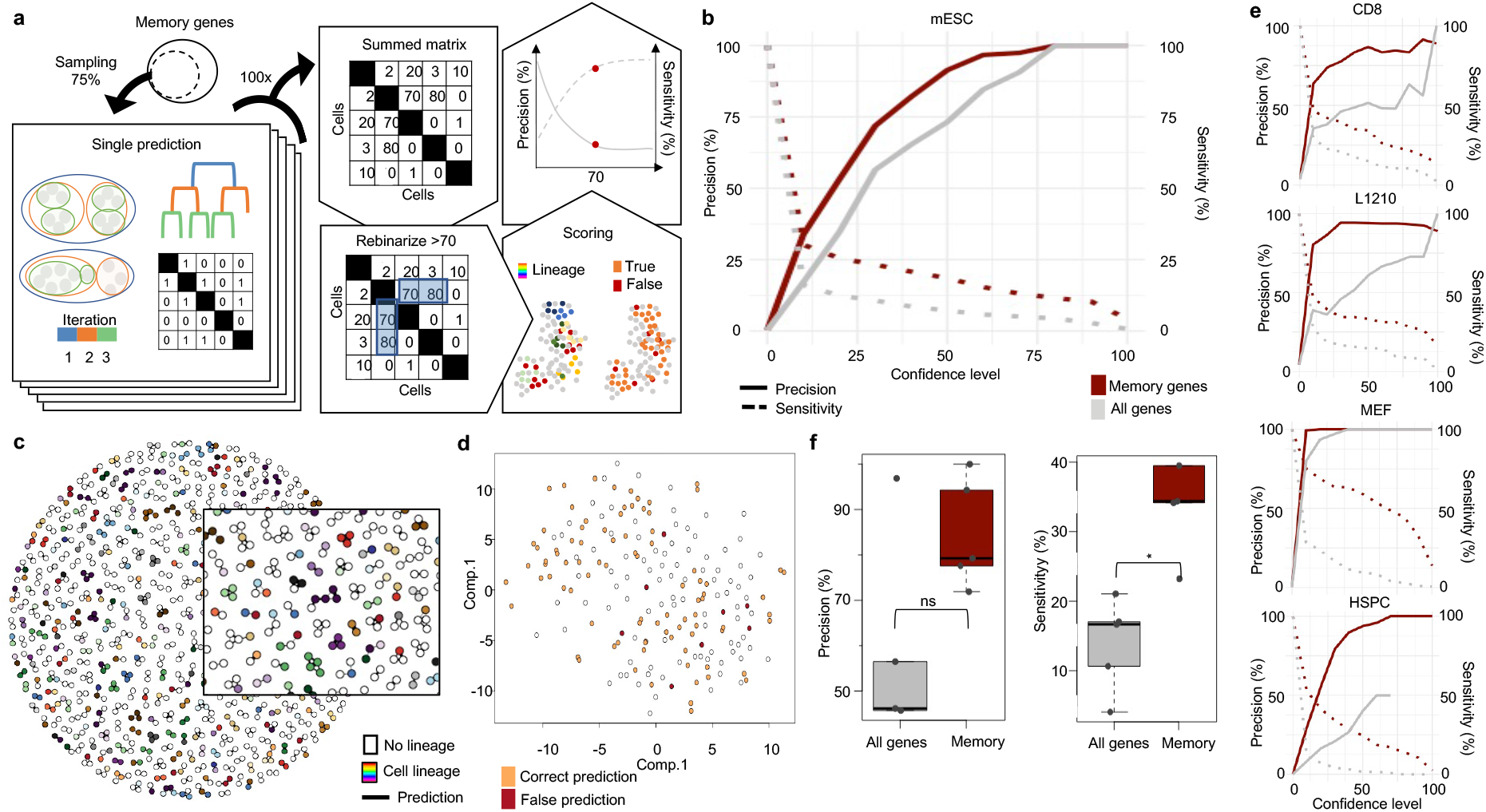
Memory genes allow to predict cell lineages. a) Scheme of the iterative hierarchical clustering algorithm developed to predict cell lineages, and of the scoring of predictions at different confidence levels using repetitive predictions. b) Curve of precision and sensitivity of lineage predictions in the mESC dataset at different levels of confidence. c) Representation of predicted lineages (black lines) with respect to the true lineages at confidence level 30. Inlet is a higher magnification. d) tSNE embedding on a PCA of the mESC dataset on all genes with coloring of cells according to tthe accuracy of predictions at confidence level 30. Selection on cell lineages of 3 members for ease of visualization. e) Precision-sensitivity curve of lineage predictions as in b for different cell types. f) Precision and sensitivity for lineage predictions across all cell types based on all genes or memory genes at confidence level 30. Boxes: intervals between the 25th and 75th percentile and median (horizontal line). Error bars: 1.5-fold the interquartile range or the closest data point when no data point is outside this range. For f significance was tested using a Mann-Withney U test (p= 0.05).

Given the overlap in memory genes across cell types, we next asked whether lineages could be predicted across datasets of the same and different cell types using a given cell type-specific set of memory genes, or using memory genes shared between several cell types. To test this, we used the previously described datasets and additional HSPC datasets (Supplementary Table 1), corresponding to different datasets from the same experiment and from different experiments in the same or related cell types. As expected, memory genes had a higher sharing across HSPC datasets than across datasets from different cell types (Extended Data Fig.5a-b). Predictions using memory genes were more precise than using all genes in the same and related cell types (Extended Data Fig.5c-d). Predictions using memory genes from other cell types or shared between several cell types performed similarly to all genes (Extended Data Fig.5c-f). Therefore, memory genes identified in one cell type can improve lineage predictions across datasets of the same and related cell types.

### Shared memory gene characteristics allow de novo lineage predictions from scRNA-seq datasets

As the expression distributions of memory genes were highly conserved across cell types, we next tried to identify a gene expression signature specific to memory genes and use it to select genes for de novo cell lineage prediction. Interestingly, memory genes were highly enriched when selecting genes on a high expression mean (quantitative memory gene characteristic) and high variability (mean-corrected CV^2^; qualitative memory gene characteristic; Fig.4a-b, Extended Data Fig.6a, see Methods). This allowed us to predict cell lineages with precision and sensitivity reaching on average 79%+/-14% and 22%+/-12%, respectively at a confidence level of 50 (Fig.4c-e, Extended Data Fig.6b-d for other confidence levels and controls). De novo lineage predictions also performed well on two additional lineage-annotated scRNA-seq data series of WM989 melanoma cells and HSCs that were not used to call memory genes (^9,27^, Table 1, and Supplementary Table 1, Fig.4c-d, Extended Data Fig.6e for other datasets). Training a neural network for gene selection did not further enrich memory genes or improve prediction values (see Methods, Extended Data Fig.6f-h). Changing the prediction parameters, such as the fraction of genes sampled for each clustering, can increase precision at the expense of sensitivity (see Methods; Extended Data Fig.7a-h). In summary, selecting genes based on expression mean and variability allows memory gene enrichment and de novo prediction of cell lineages in scRNA-seq datasets. We named this approach <u>G</u>ene <u>E</u>xpression <u>M</u>emory-based <u>L</u>ineage Inference or GEMLI.

**Figure 4:**
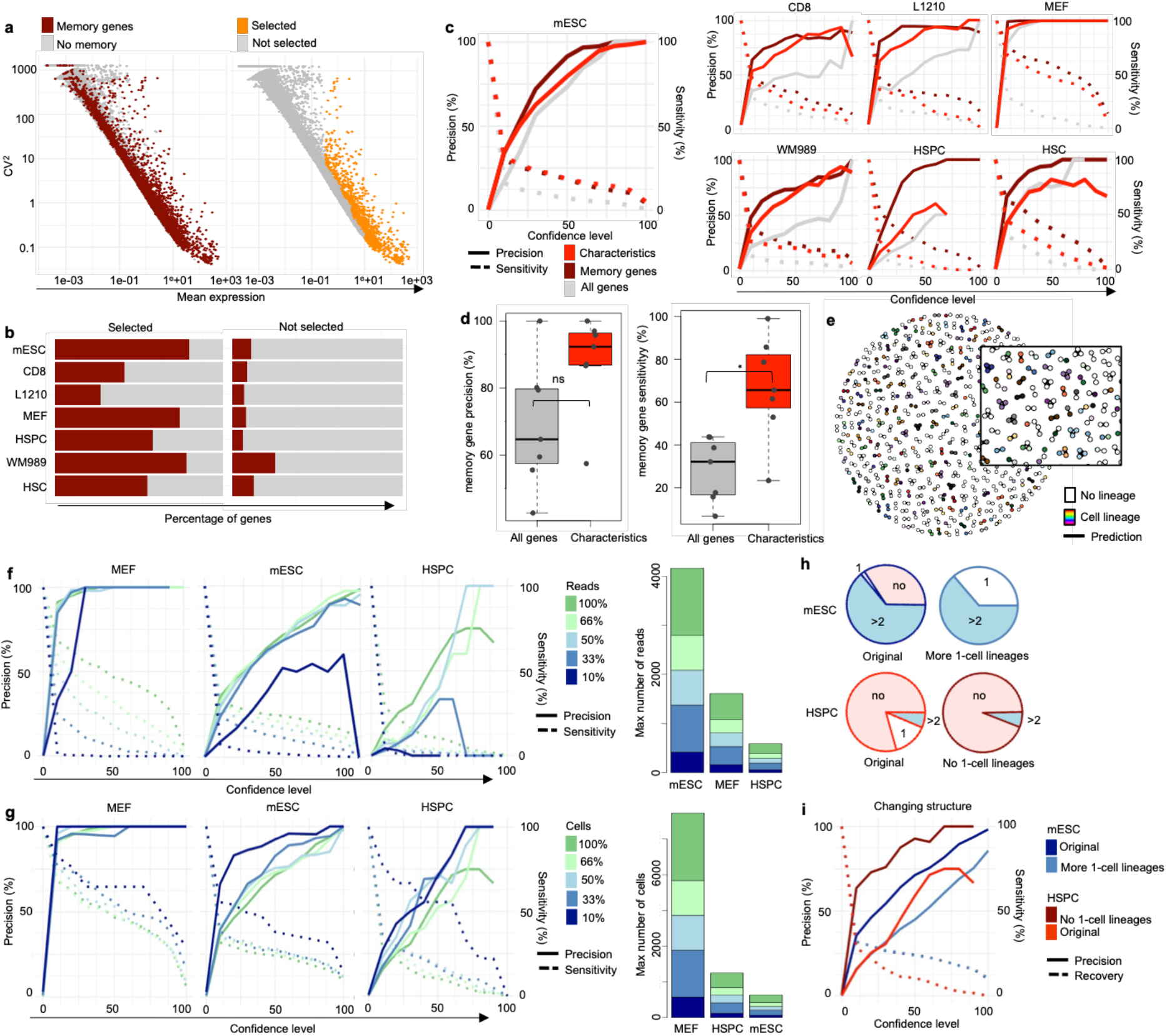
Lineage relations can be predicted de novo across cell types. a) Scatterplots of mean expression against mean-corrected CV_2_ highlighting memory genes (left) or characteristics-based selected genes (right). b) Percentage of memory genes in the selected and non-selected genes with memory-gene characteristics. c) Precision-sensitivity curves of lineage predictions in different cell types for memory genes, all genes, and genes selected based on memory-gene characteristics. d) Percentage of precision (top) and sensitivity (bottom) of memory-gene based lineage predictions gained using all genes or genes chosen based on memory-gene characteristics at confidence level 50. Boxes: intervals between the 25th and 75th percentile and median (horizontal line). Error bars: 1.5-fold the interquartile range or the closest data point when no data point is outside this range. Significance was tested using a Mann-Withney U test (p= 0.05). e) Representation of predicted lineages (black lines) with respect to the true lineages at confidence level 50. Inlet is a higher magnification. f) Precision-sensitivity curves for the MEF, mESC, and HSPC dataset after subsampling reads as indicated. The bar graph represents the max read number in the indicated dataset and subsampling category. g) Same representation as in f for subsampling of cells. h) Representation of the mESC and HSPC datasets showing the fraction of cells in lineages of the indicated sizes before and after changing the lineage structure. no: cells without an assigned lineage. i) Precision-sensitivity curves for the mESC and HSPC datasets after changing their lineage structure as indicated in h.

### GEMLI performance depends on sequencing depth, size and lineage structure

As precision and sensitivity of GEMLI varied between datasets, we asked whether this could be explained by underlying differences in cell lineage structure, size, or sequencing depth (Supplementary table 1, Fig.4f-i). We applied GEMLI to the mESC, MEF and/or HSPC dataset, after downsampling reads, cells, or changing lineage structure (see Methods). For mESCs, which had the highest sequencing depth, lower read counts led to both lower precision and sensitivity. This suggests that high sequencing depth is critical for high-quality predictions (Fig.4f). Subsamples with fewer cells generally had a slightly better precision and sensitivity, especially for very low read counts (Fig.4g). We next focused on the impact of single-member lineages on predictions, which can occur through differences in the experimental setup or because of technical artifacts of barcode-based lineage assignment (see Methods). Many single member lineages were present in the HSPC data, and few in the mESCs dataset (Fig.4h). Including cells without barcode assignment as single member lineages in the mESC dataset lowered precision, while excluding them in the HSPC data had the opposite effect (Fig.4i). In summary, differences in the size, lineage structure or assignment, and sequencing depth of scRNA-seq datasets can partly explain differences in prediction performance.

### GEMLI predicts cell lineages after long culture times and assigns cells to individual crypts and organoids

Next, we wondered if cell lineages could be predicted over long culture times. Several studies have reported gene expression memory above 5 cell divisions ^8,10,20,34^. In line with this, the over-time point gene expression correlation in the 28-day MEF reprogramming time course (Supplementary table 1) was higher for related than for random cell pairs up to day 15 (Extended Data Fig.8a). GEMLI stayed highly precise for all datasets, however sensitivity decreased over culture and replating time (Fig.5a, Extended Data Fig.8b, see Methods). We next analyzed scRNA-seq data of intestinal organoids grown for up to 6 days and crypts from adult mice, which have a turnover of around 6 days, and for which the structure (crypt or organoid) of origin is known for each cell (^32^, Table 1, Supplementary table 1, see Methods). De novo predictions of small cell lineages grouped cells of the same origin with very high precision (Fig.5b-d, Extended Data Fig.8e). Interestingly, here also larger cell groups could be assigned to single organoids or crypts correctly (Fig.5d-f Extended Data Fig.8f-h), which allowed to analyze the variability in individual crypts and organoid composition (Fig.5g, Extended Data Fig.8i). To summarize, GEMLI remains highly precise on long culture times, and distantly related cells can be identified accurately enough to define individual small structures such as intestinal organoids and crypts.

**Figure 5:**
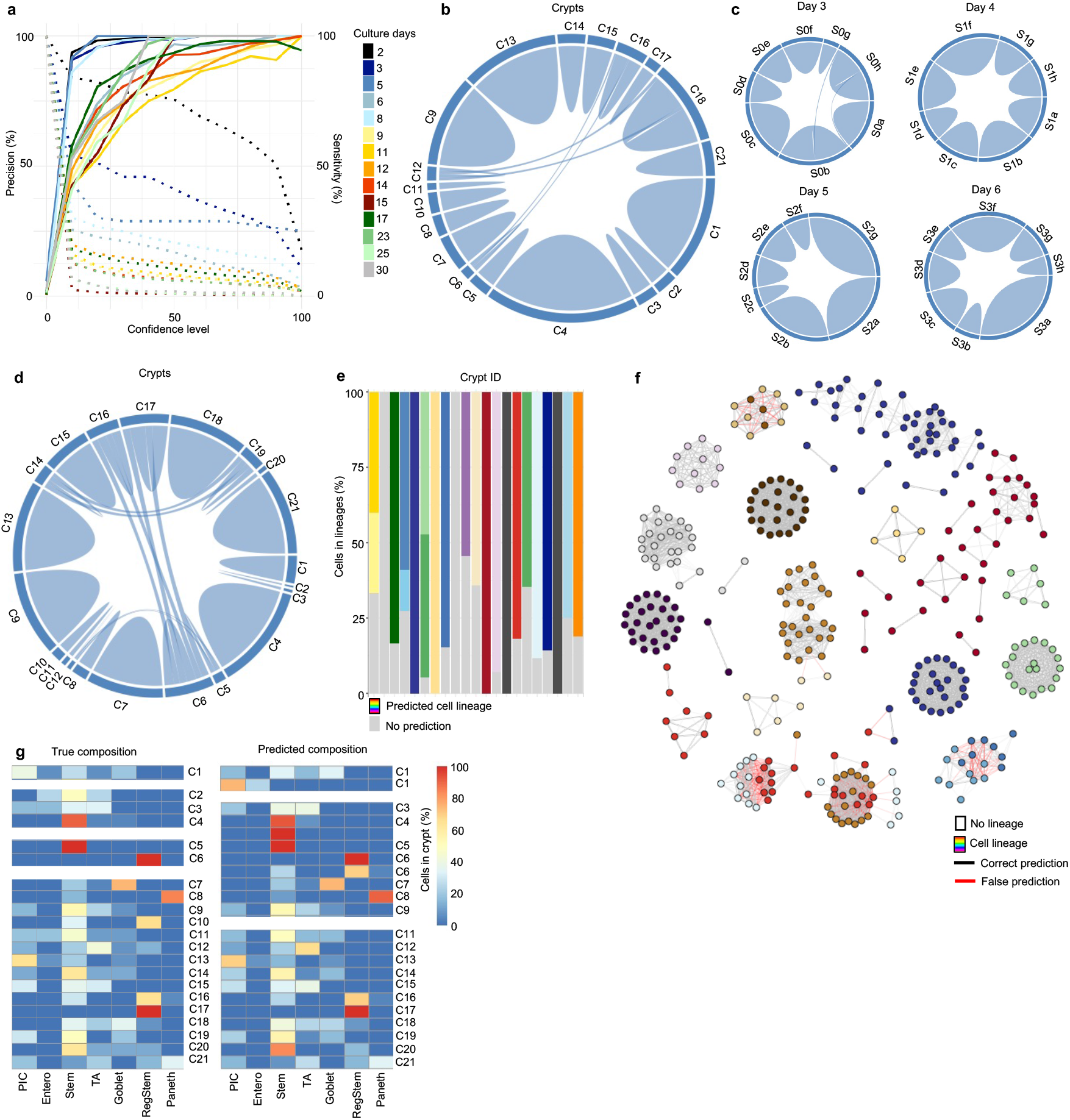
De novo lineage predictions over extended culture times correctly assign cells to individual crypts and organoids. a) Precision-sensitivity curves of de novo lineage predictions for cell lineages for MEF datasets from different culture timepoints (n=4 for day 2, 8 and 9, and n=2 for all other culture timepoints). Mean is represented. b) Chord diagram showing predicted lineages at confidence level 50 with their relation to the individual intestinal crypts (dark boxes at outer circle). Lines starting and ending in the same dark box are correct predictions. Lines crossing the circle are false predictions. c) Chord diagram as in b for intestinal organoids split by culture timepoints. d) Chord diagram as in b for predictions allowing lineages of 5-40 cells in intestinal crypts. e) The percentage of cells attributed to individual lineages for each crypt when predicting lineages of 5-40 cells. f) Representation of crypt predictions at confidence level 70). Color of connection lines indicates accuracy of predictions. g) The percentage of cells belonging to the indicated cell type for individual crypts (left) and for predicted lineages of 5-40 cells (right). For the predicted lineages, the crypt of origin is indicated. PIC, potential intermediate cluster; Entero, enterocyte cluster; Stem, stem cells; TA, transit amplifying cells; RegStem, regenerative stem cells.

### GEMLI identifies memory gene categories, cell fate decisions and associated gene expression programs

We next tested the performance of GEMLI in analyses previously performed on experimentally lineage-annotated scRNA-seq datasets. We compared memory genes and associated GO-terms from GEMLI predictions and true lineages, and found these to be highly correlated for all datasets (Fig.6a, Supplementary table 5, Extended Data Fig.9a-c). Next, we analyzed if GEMLI can inform on cell fate choices. Both in WM989 melanoma cell and HSPC datasets (total of 56 datasets), the fraction of related cells among different symmetric and asymmetric phenotype pairs (For WM989 cells, BRAF- and MEK-susceptible, as well as drug-resistant phenotypes, for HSPC 10 different hematopoietic phenotypes, see Methods) correlated well for GEMLI predictions and barcode lineages (Extended Data Fig.9g-h, Fig.6b). Including phenotype pairs predicted on cells without recovered barcodes, led to a similar correlation, and increased the number of analyzed cell pairs to 38%+/-23 and 339%+/-236 of the number of barcode lineages for HSPCs and WM989 cells respectively (Extended Data Fig.9i-k). Differential expressed genes (DEG) between asymmetric phenotype pairs and symmetric phenotype pairs were highly correlated between barcode and GEMLI predictions in both cell types (Fig.6c-d). DEG analysis between cell pairs of undifferentiated and neutrophil cells in the HSPC data recovered genes associated with the regulation of early neutrophil differentiation such as *Wfdc21, Ltf, Lyz2, Lcn2, Ngp* and *Camp* ^35^ in both predicted and barcode lineages (Fig.6e). In summary, GEMLI allows to predict diverging cell fate choices and to analyze gene expression programs of cells switching to a particular fate solely from scRNA-seq datasets, and can have higher recoveries of lineages than barcode-based assignments.

**Figure 6:**
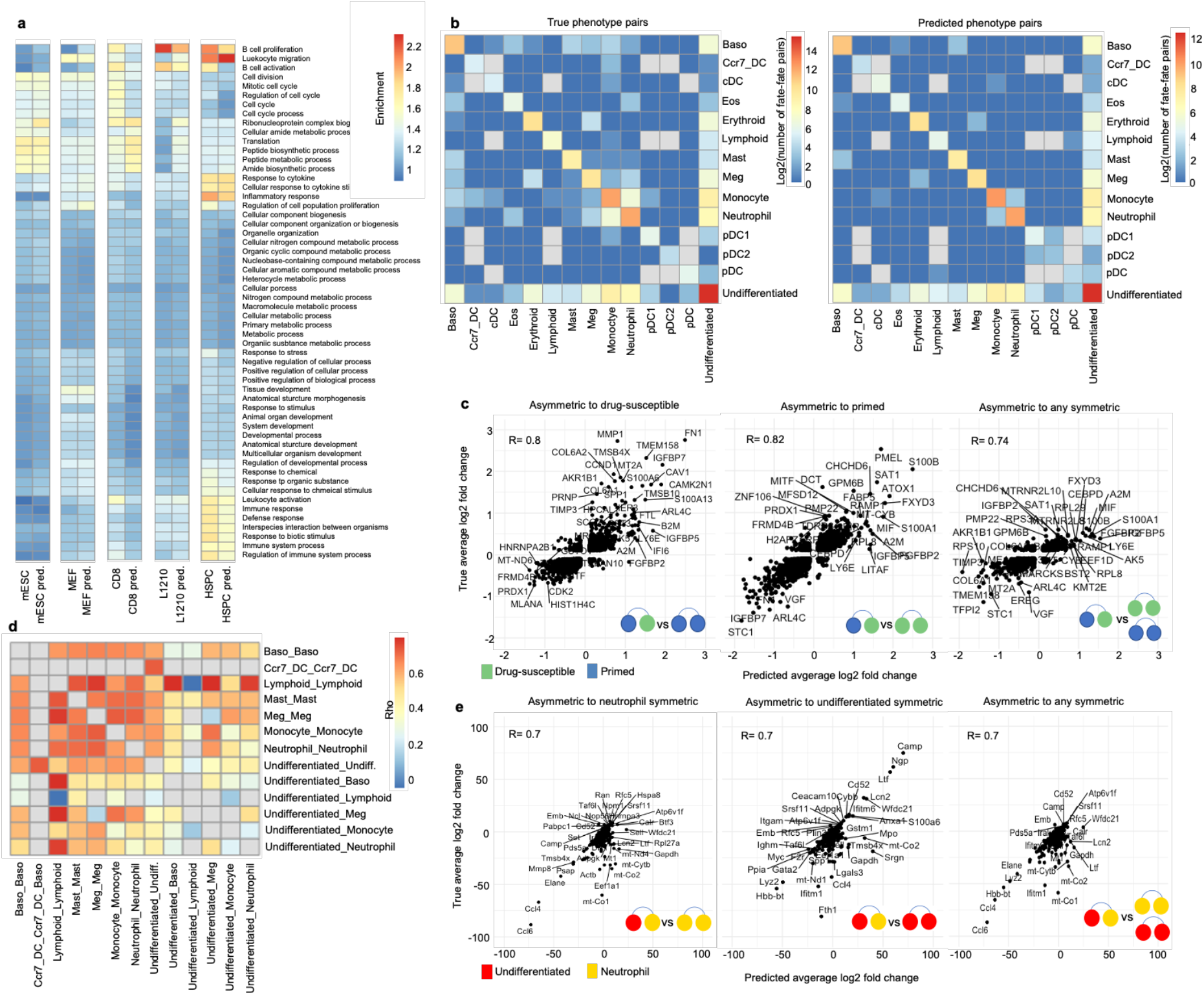
De novo lineage predictions allow retrieving cell-type specific memory GO terms and to quantify diverging cell-fate decisions. a) Heatmap of combined top 10 GO-term enrichment in memory genes called on true or de novo predicted lineages at confidence level 30 in different cell types. Heatmap of the number of cell pairs (log) in true (left) and de novo predicted (right) cell lineages in all possible phenotype pair categories in the 48h and 96h HSPC datasets (n=48). The sum across all datasets is shown. c) Scatterplot of the average log2 fold change for DEG called between true and de novo predicted entirely drug susceptible (left), entirely primed cell lineages (middle), or both (right), and cell lineages with members in both states (asymmetric cell pairs) across all WM989 datasets (n=8). Each dot represents one DEG in one of the datasets. Spearman rank correlation is given. The top 20 and top 10 of highest and lowest enriched genes in asymmetric cell pairs are named. d) Heatmap of the Spearman correlation in DEG between cell pairs in the indicated phenotype categories across HSPC datasets as in b. Grey are categories for which no DEG could be called. Phenotype pair categories for which no DEG could be called in any of the datasets are omitted. e) Scatterplot of the average log2 fold change for DEG called between true and de novo predicted entirely neutrophil (left), entirely undifferentiated (middle), or both (right), and cell lineages with members in both states (asymmetric cell pairs of undifferentiated cells and neutrophils) across one HSPC dataset. Each dot represents one DEG in one of the datasets. Spearman rank correlation is given. The top 20 and top 10 of highest and lowest enriched genes in asymmetric cell pairs are named. For b and d: Baso, basophil; Eos, eosinophils; Meg, megakaryocytes; pDC1/2, plasmacytoid dendritic cells 1/2; Ccr7+_DC, migratory dendritic cells; cDC, classic dendritic cell.

## Discussion

We describe GEMLI, a computational pipeline that allows de novo prediction of cell lineages from scRNA-seq datasets based on the identification of genes that maintain their expression level through cell division. GEMLI is applicable to scRNA-seq datasets without lineage information, thereby alleviating the need for experimental procedures to assign lineages before sequencing. GEMLI is available as a R package at GitHub (https://github.com/UPSUTER/GEMLI) and encompasses functions for de novo cell lineage predictions and memory gene identification. GEMLI is immediately applicable to existing scRNA-seq datasets generated by various technologies. As for any quantitative analysis, sequencing depth influences its performance, but based on subsampling analysis, we expect predictions to be reliable at sequencing depths above 5,000-8,000 reads/cell, as commonly achieved for available scRNA-seq technologies ^36–38^. Another limitation stems from the structure of the dataset, such as when only a small fraction of cells from a large population is sequenced, resulting in a low number of closely related cells.

Other methods to infer cell lineages from scRNAseq data alone have been reported but operate on different timescales. Lineage inference from single nucleotide polymorphisms, copy number variations, or mitochondrial mutations recovered in scRNA-seq data ^39–43^ can identify lineages extending over long time spans (weeks to months) and requires high coverage (long reads) and sequencing depths. At the other end of the spectrum, transcriptome trajectory inference algorithms, although not inferring lineages, can predict gene expression changes on the order of hours, using exonic reads alone ^44^, or RNA velocity based on exonic and intronic reads ^33,45^. GEMLI is ideally suited for intermediate time spans, i.e. to analyze small cell lineages and gene expression changes over the course of a few cell divisions (days), and requires only exonic reads. It could be used in conjunction with orthogonal lineage tracing or inference approaches that identify lineages on other timespans to refine lineage pedigrees.

GEMLI allows to predict cell lineages but also reliably identifies genes displaying gene expression memory, thus opening the door to understanding underlying epigenetic mechanisms in different cell types. The identification of different types of cell divisions (symmetric or asymmetric) by GEMLI can be used to identify cell fate regulators directly from scRNA-seq datasets, alleviating the need for experimental lineage annotation that has been used so far in this context ^9,17,19,27^. Notably, GEMLI-predicted lineages can be used as input for trajectory inference and cell fate decision analysis designed for lineage-annotated scRNA-seq data ^46–48^. In datasets that are lineage-annotated through cellular barcodes, GEMLI can increase the recovery of lineages through the predictions of lineages for cells without barcode expression or not meeting other criteria of barcode-based lineage identification. Finally, the ability of GEMLI to assign individual cells to their larger structure of origin could be applicable not only to organoids or crypts as shown here, but also to other structures emerging from a common ancestor or a small pool of stem cells, such as glands, spatially restricted areas of the skin or small metastases. GEMLI should thus become a method of choice in situations where orthogonal lineage identification approaches are not feasible, and may serve as a powerful discovery tool using already existing scRNA-seq datasets.

## Extended Data Figures

**Extended Data Figure 1:**
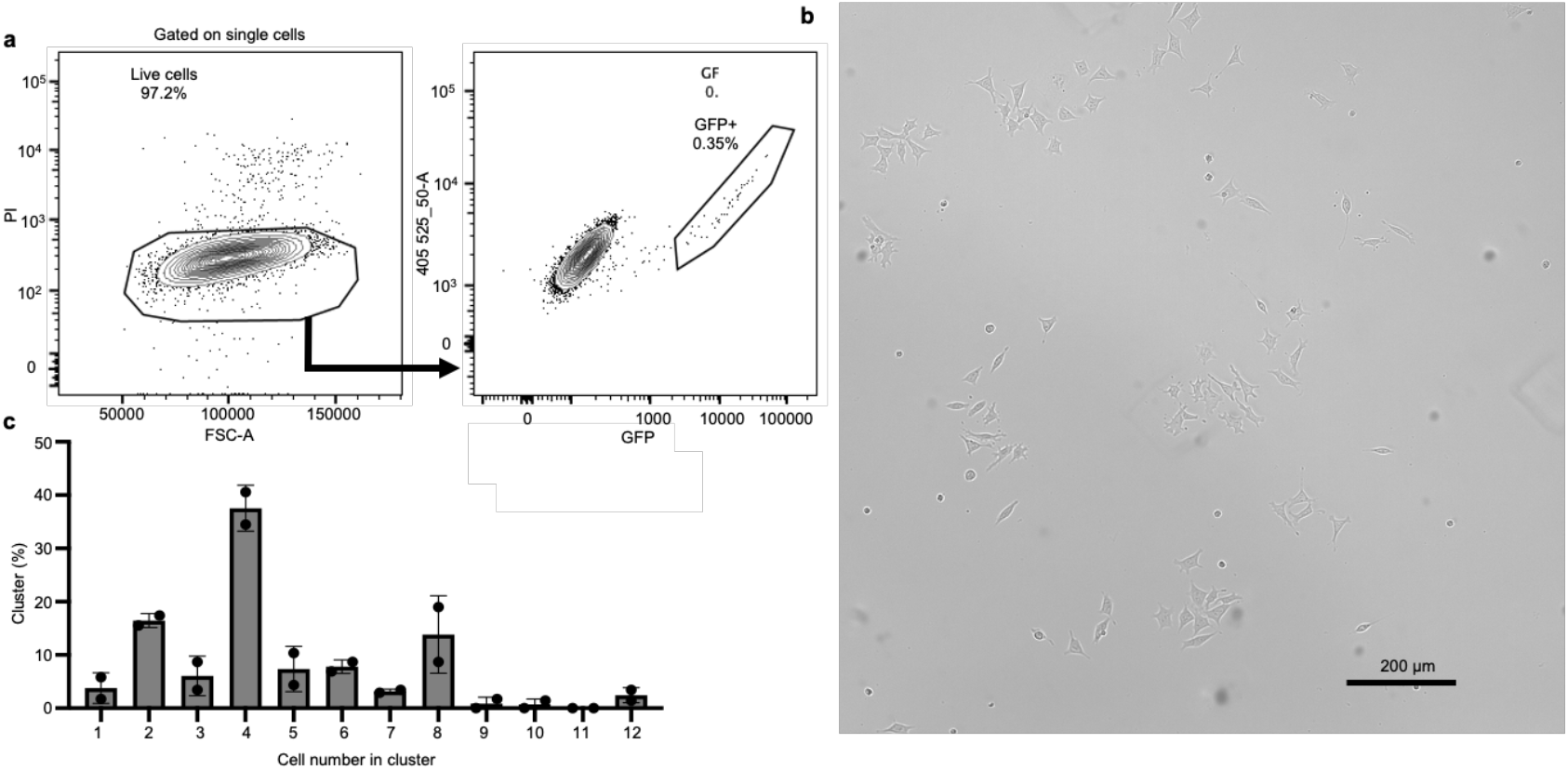
Generation of a mESC lineage-annotated scRNA-seq dataset. a) Barcoded cells were sorted as single, live GFP_+_ cells into two wells of a 96-well plate. b) Brightfield image of mESCs after 48h of culture. c) Distribution of cluster sizes of mESCs after 48h of culture (mean of two wells) counted manually in microscopy images. The low seeding density in this experiment allowed related cells to remain clustered.

**Extended Data Figure 2:**
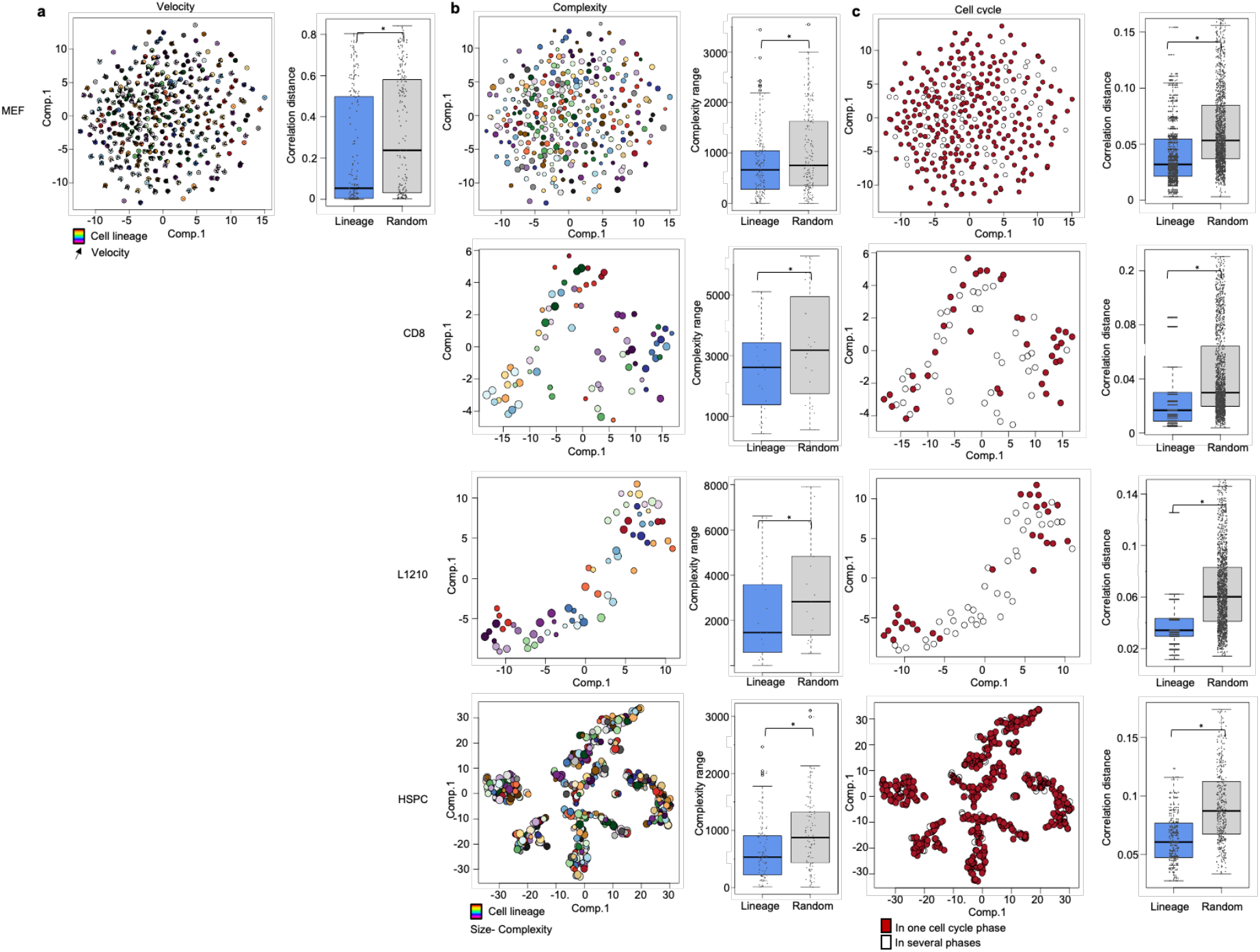
Gene expression memory in scRNA-seq data across cell types. a) tSNE embedding of a PCA on exonic and intronic data of the lineage-annotated (colors) MEF datasets including velocity vectors (arrows, left) and the comparison of log (correlation distance) for cells belonging to the same cell lineage (blue), and randomly sampled cells (gray) in the same dataset. Boxes: intervals between the 25th and 75th percentile and median (horizontal line). Error bars: 1.5-fold the interquartile range or the closest data point when no data point is outside this range. b-c) tSNE embedding of a PCA on exonic data of the lineage-annotated CD8, L1210, and HSPC, and on exonic and intronic data of the MEF datasets as in a, size as complexity quantile (b), or coloring according to the completeness of a lineage in one cyclone-assigned cell-cycle phase (c). Boxplots: respective quantification of similarity in related and randomly sampled cells (as correlation distance or complexity range). Boxplot as in a. For ease of visualization, cell lineages of 2 (MEF), or 1-5 members (all other datasets) are selected for the embeddings. Statistical significance for (a-c) was tested using a Mann-Whitney U-test (p= 0.05).

**Extended Data Figure 3:**
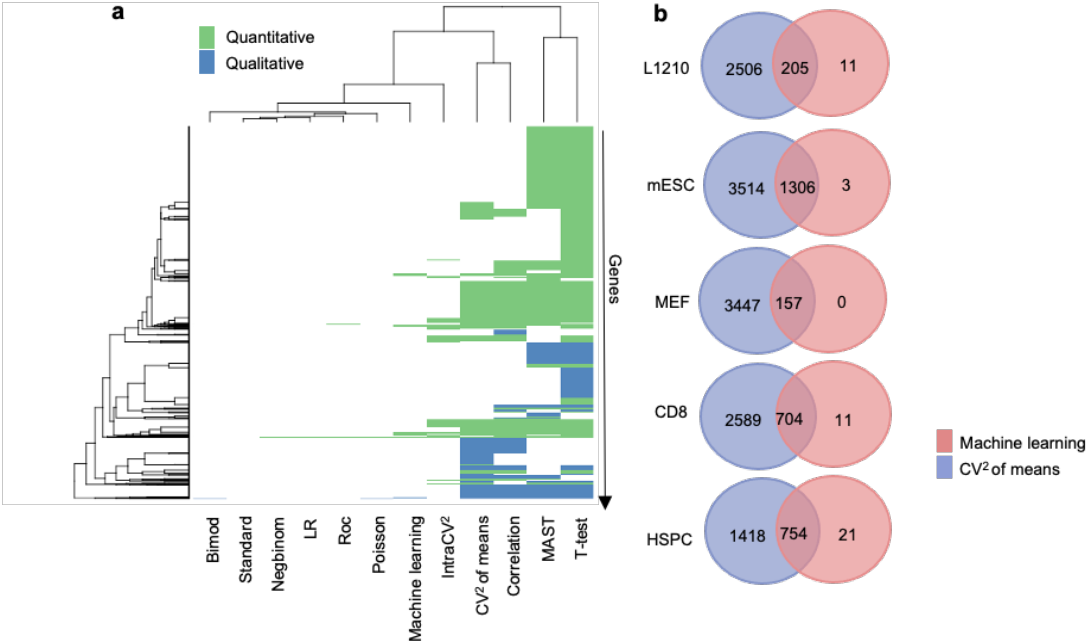
Overlap of memory genes identified using different criteria and methods. a) Heatmap showing the sharing of memory genes called using different methods. Red: quantitative memory genes; Green: qualitative memory genes. Memory genes are selected based on a high correlation of gene expression within cell lineages (correlation), a small intra-cell lineage variability (intraCV_2_), a large variability across means of cell lineages (CV_2_ of means), are marker genes for cell lineages found using Seurats FindMarker() function, or have been selected using a machine learning approach (see Methods). b) Overlap in the Machine learning generated memory gene sets (red) and the CV_2_ of means-based memory gene sets (blue) of the indicated cell types.

**Extended Data Figure 4:**
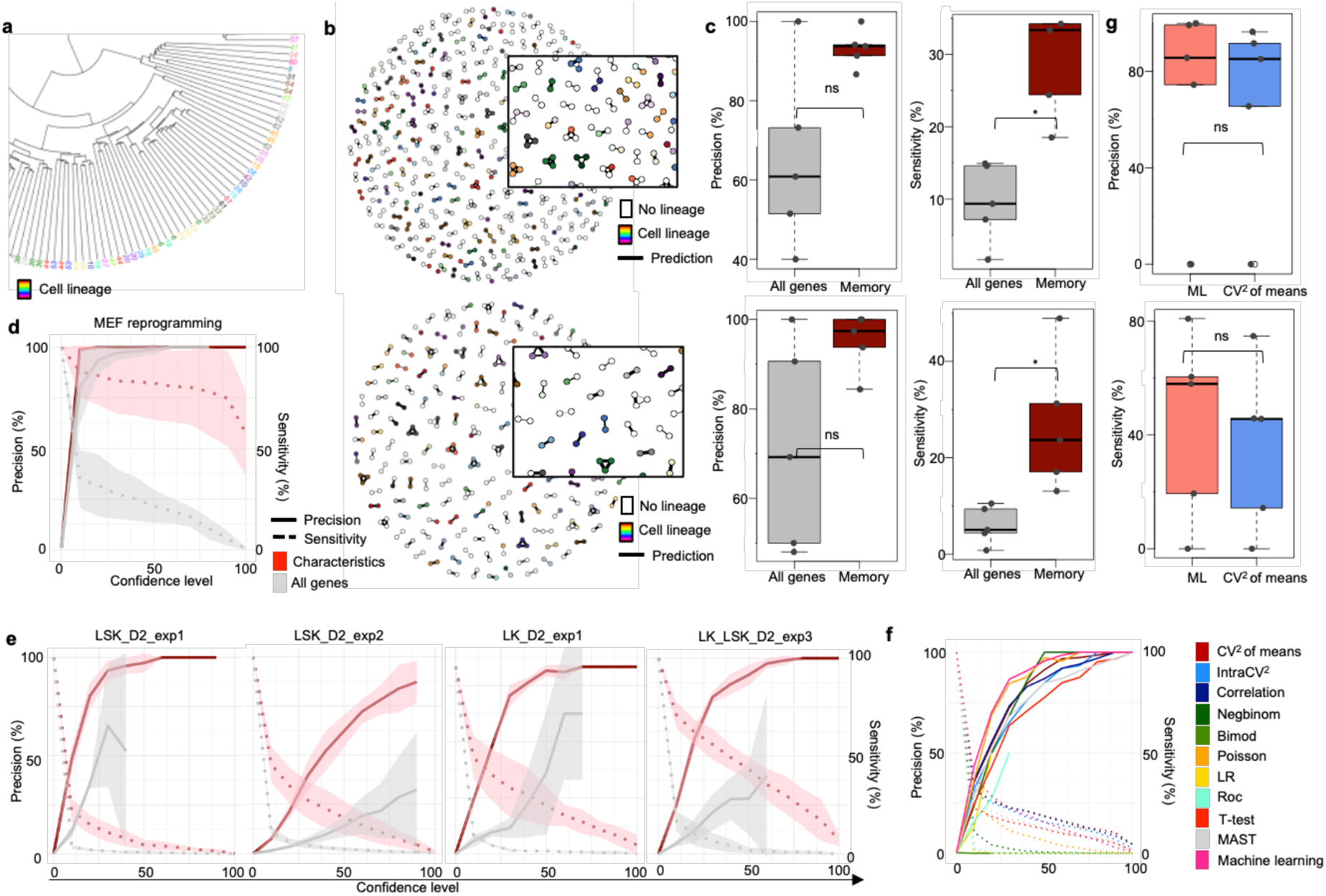
Memory genes allow for lineage predictions within the dataset of origin. Representation of a simple hierarchical clustering based on the Spearman correlation distance of the mESC scRNA-seq data. Coloring and tip number correspond to individual cell lineages. Selection on a quarter of the cell lineages of 3 members for ease of visualization. b) Representation of lineage predictions on the mESC dataset at confidence levels 50 (top), or 70 (bottom). Inlet is a higher magnification. c) Precision (left) and sensitivity (right) for lineage predictions across all cell types based on all genes or memory genes for confidence level 50 (top), or 70 (bottom). Boxes: intervals between the 25th and 75th percentile and median (horizontal line). Error bars: 1.5-fold the interquartile range or the closest data point when no data point is outside this range. d) Precision-sensitivity curves of lineage predictions in the MEF datasets (n=5) of 48h-72h culture times across the reprogramming time course. Line: mean; Shades: S.D.. e) Same representation as in D for the HSPC datasets of the experiments LSK_D2_exp1 (n=3), LSK_D2_exp2 (n=10), LK_D2_exp1 (n=3) and LK_LSK_D2_exp3 (n=6). f) Precision-sensitivity curve of lineage predictions in the mESC dataset using memory genes called as in Extended Data Fig.3 a (see Methods). The Standard gene set did not lead to any prediction and is omitted from the figure. g) Precision (left) and sensitivity (right) for predictions at confidence level 50 using memory genes identified using the machine learning approach or CV_2_ of means as indicated. Boxplot as in c. For d and h significance was tested using a Mann-Withney U test (p= 0.05).

**Extended Data Figure 5:**
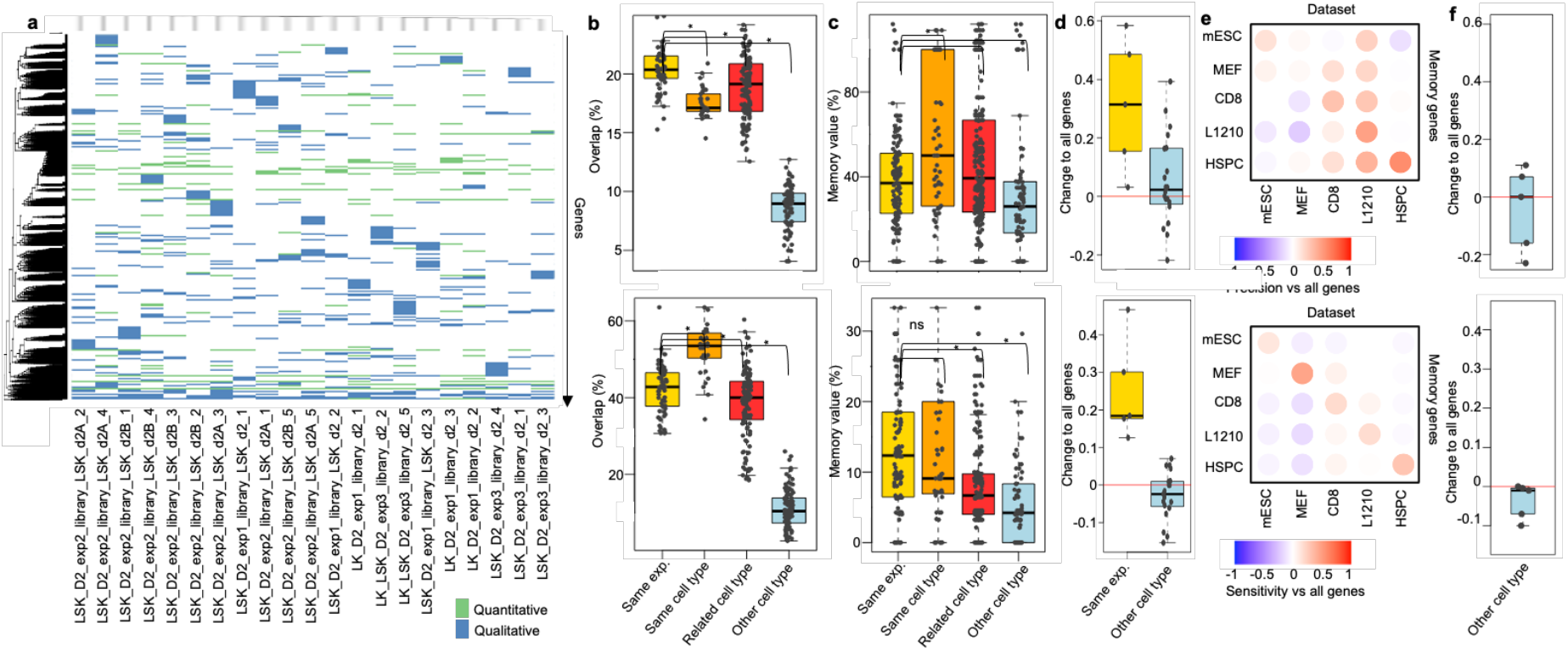
Overlap of memory genes and performance in lineage predictions across datasets and cell types. a) Heatmap showing the sharing of memory genes across HSPC datasets of the same or similar cell types. Red: Quantitative memory genes; Green: Qualitative memory genes. b) Percentage overlap in qualitative (top) or quantitative (bottom) memory genes across pairs of datasets from the same experiment (yellow; LK_D2_exp1, LK_LSK_D2_exp3, LSK_D2_exp1, and LSK_D2_exp2 - all libraries respectively), same cell type (orange; LSK - all libraries of LSK_D2_exp1 and LSK_D2_exp2), related cell type (red; all combinations of all libraries of LK_LSK_D2_exp3, LSK_D2_exp1, LSK_D2_exp2, and LK_D2_exp1) or other cell types (blue; combinations of all libraries of LK_LSK_D2_exp3, LSK_D2_exp1, LSK_D2_exp2, and LK_D2_exp1 with the datasets CD8, L1210, MEF, and mESC as in the main figures). Percentage overlap was calculated as percentage of the mean memory gene number of the two compared datasets. Boxes: intervals between the 25th and 75th percentile and median (horizontal line). Error bars: 1.5-fold the interquartile range or the closest data point when no data point is outside this range. c) Percentage of memory gene value in precision (top) and sensitivity (bottom) at confidence level 50 for the categories indicated as in b. Boxplot as in b. d) Difference in precision (top) and sensitivity (bottom) at confidence level 30 for memory genes of the dataset itself or of other cell types as in b compared to all genes. Line in red depicts no change to all genes. Boxplot as in b. e) Heatmap showing the difference of precision (top) and sensitivity (bottom) of lineage predictions at confidence level 30 based on memory genes vs all genes across cell types. f) Difference in precision (top) and sensitivity (bottom) at confidence level 50 for predictions using memory genes shared in four dataset to predict in the fifth compared to predictions using all genes. Line in red depicts no change to all genes. Boxplot as in b. For b-c significance was tested using a Mann-Withney U test (p= 0.05).

**Extended Data Figure 6:**
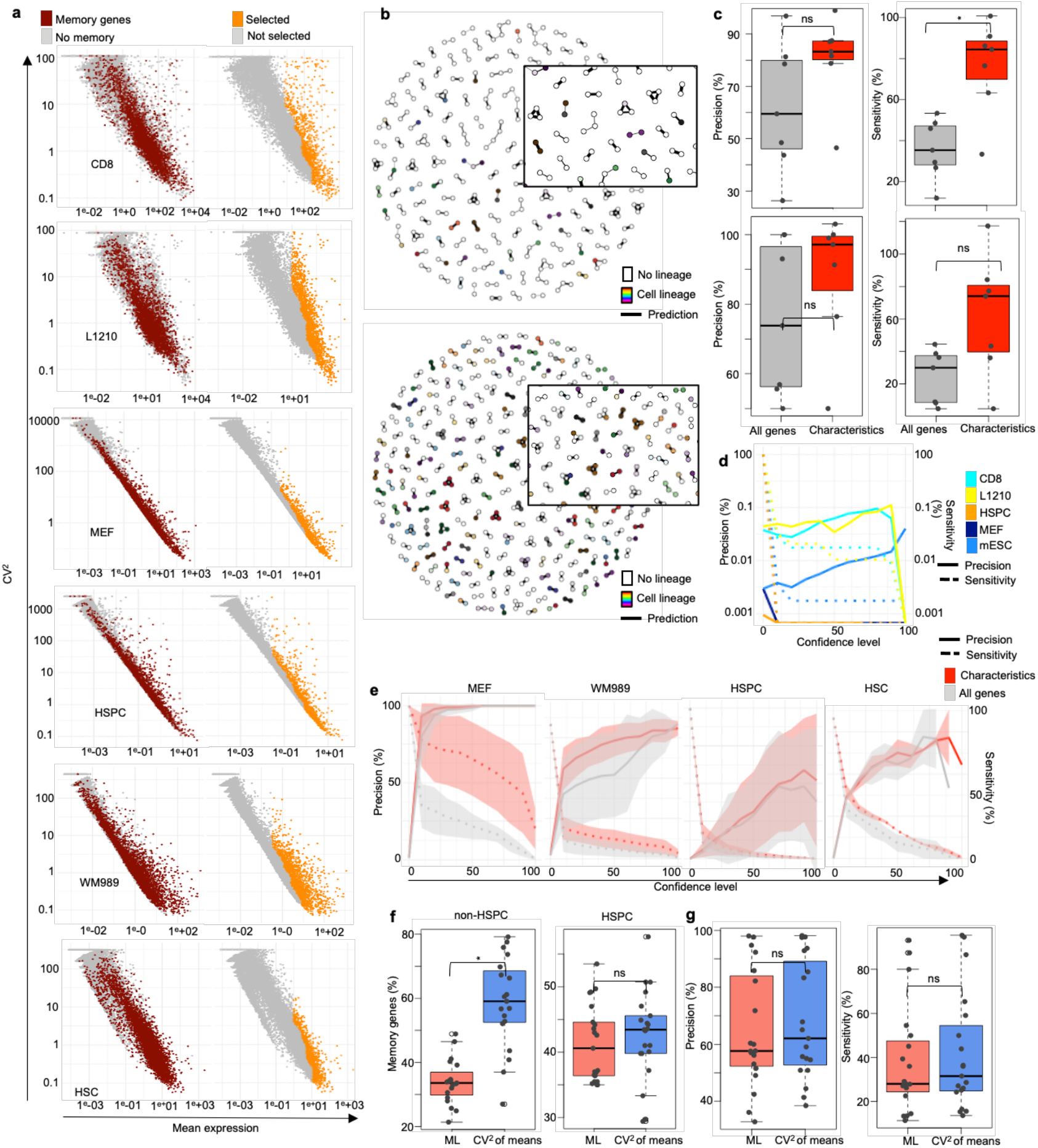
De novo lineage predictions based on genes with memory gene characteristics. a) Scatterplots of mean expression against mean-corrected CV_2_ highlighting memory genes (left) or characteristics-based selected genes (right). b) Representation of the lineage predictions on the mESC dataset for confidence levels 30 (top) and 70 (bottom). For confidence level 30 only half of the cells are represented for ease of visualization. Inlet is a higher magnification. c) Boxplot of precision (left) and sensitivity (right) for lineage predictions across all cell types based on all genes or characteristics selected genes for confidence level 50 (top), or 70 (bottom). Boxes: intervals between the 25th and 75th percentile and median (horizontal line). Error bars: 1.5-fold the interquartile range or the closest data point when no data point is outside this range. d) Precision-sensitivity curves of de novo lineage predictions of random cell lineages for the indicated datasets. e) Precision-sensitivity curves of lineage predictions in the following datasets: MEF (n=5) of 48h-72h culture times across the reprogramming time course, HSPC (LK, LSK, and LK_LSK mix, n=20, HSC (n=2), and WM989 (n=8). Line: mean; Shades: S.D.. f) Percentage of memory genes when selected based on memory gene characteristics or a machine learning approach in all datasets of 48h or 96h across mESC, MEF, WM989, HSC (non-HSPC, left, n=19) or HSPC (right, n=21). Boxplot as in c. g) Precision (left) and sensitivity (right) for all non-HSPC datasets as in F at a confidence level of 50 in the python prediction implementation. Boxplot as in c. For HSPC datasets no lineages were predicted. For c, f and g, significance was tested using a Mann-Withney U test (p= 0.05).

**Extended Data Figure 7:**
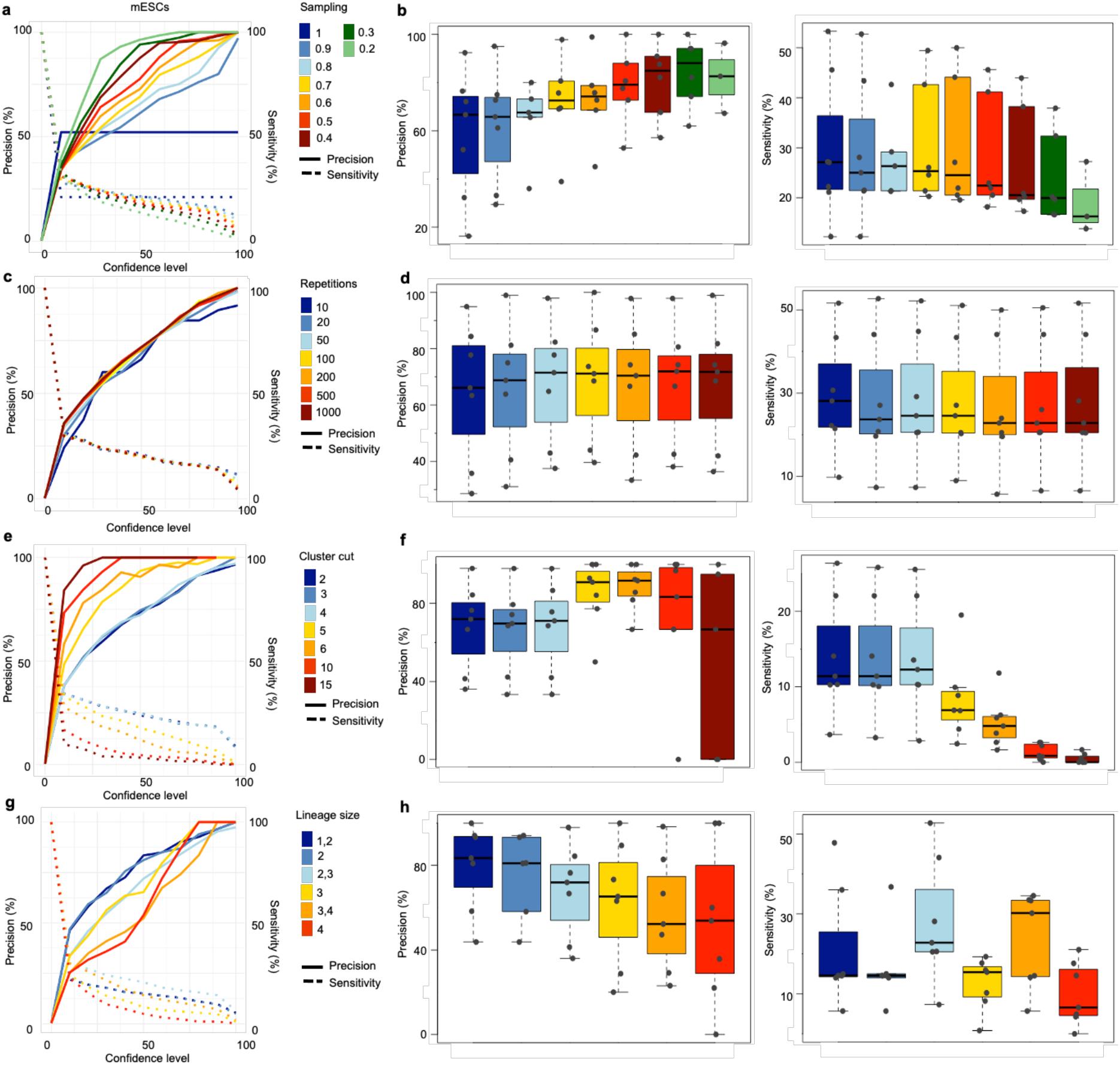
Parameters of de novo lineage predictions based on genes with memory gene characteristics. a) Precision-sensitivity curves of lineage predictions in the mESC dataset sampling different percentages of genes during each iterative clustering as indicated. b) Precision (left) and sensitivity (right) for mESC, CD8, L1210, HSPC, HSC, WM989, MEF datasets at a confidence level of 50 using different percentages of genes (color-coded) as in a. Boxes: intervals between the 25th and 75th percentile and median (horizontal line). Error bars: 1.5-fold the interquartile range or the closest data point when no data point is outside this range. c) Precision-sensitivity curves of lineage predictions in the mESC dataset repeating the iterative clustering different number of times as indicated. d) Precision and sensitivity for datasets as in b using different number of repetitions (color-coded) as in c. e) Precision-sensitivity curves of lineage predictions in the mESC dataset stopping iterative clustering at clusters with different cell numbers as indicated. f) Precision and sensitivity for datasets as in b using stopping iterative clustering at cluster sizes (color-coded) as in e. g) Precision-sensitivity curves of lineage predictions in the mESC dataset asking for different lineage sizes as indicated. h) Precision and sensitivity for datasets as in b calling lineages of different sizes (color-coded) as in e.

**Extended Data Figure 8:**
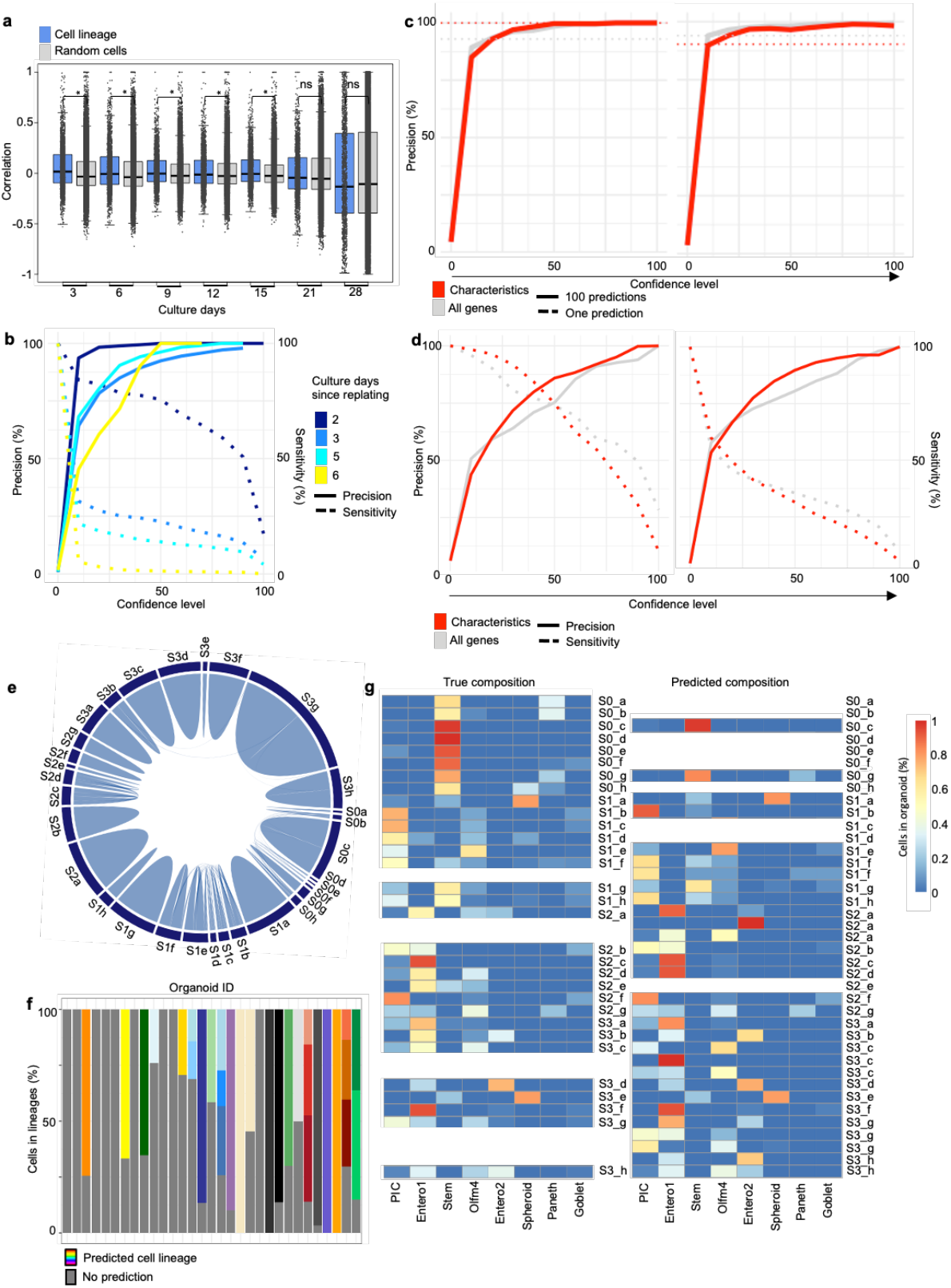
Performance of de novo lineage predictions after long culture time spans. a) Correlation in memory gene expression of day 0 for related cell pairs shared between day 0 and the indicated time points or random cell pairs across MEF datasets of 48h-72h across the reprogramming time course (MEF_2 replicate; n=3). Boxes: intervals between the 25th and 75th percentile and median (horizontal line). Error bars: 1.5-fold the interquartile range or the closest data point when no data point is outside this range. b) Precision-sensitivity curves of de novo lineage predictions for the indicated MEF datasets ordered by time since replating (n=4, 20, 6, and 6 for day 2, 3, 6, and 7 respectively). c) Precision of lineage predictions of small cell lineages in intestinal organoids (left) and crypts (right) for all genes, and genes selected based on memory gene characteristics. Dotted lines: precision of one single iterative clustering without subsampling. d) Precision-sensitivity curves for lineage predictions allowing sizes of 5-40 cells for crypts (left) and organoids (right). e) Chord diagram showing predicted lineages allowing sizes of 8-40 cells at confidence level 70 with their relation to the individual intestinal organoid (dark boxes at outer circle). Lines starting and ending in the same dark box are correct predictions. Lines crossing the circle are false predictions. f) The percentage of cells attributed to individual lineages for each organoid when predicting lineages of 8-40 cells. g) The percentage of cells belonging to the indicated cell type for individual organoids (left) and for predicted lineages of 5-40 cells (right). For the predicted lineages the organoid of origin is indicated. PIC, potential intermediate cluster; Entero1/2, enterocyte cluster1/2; Stem, stem cells. Significance in a was tested using a Mann-Withney U test (p= 0.05).

**Extended Data Figure 9:**
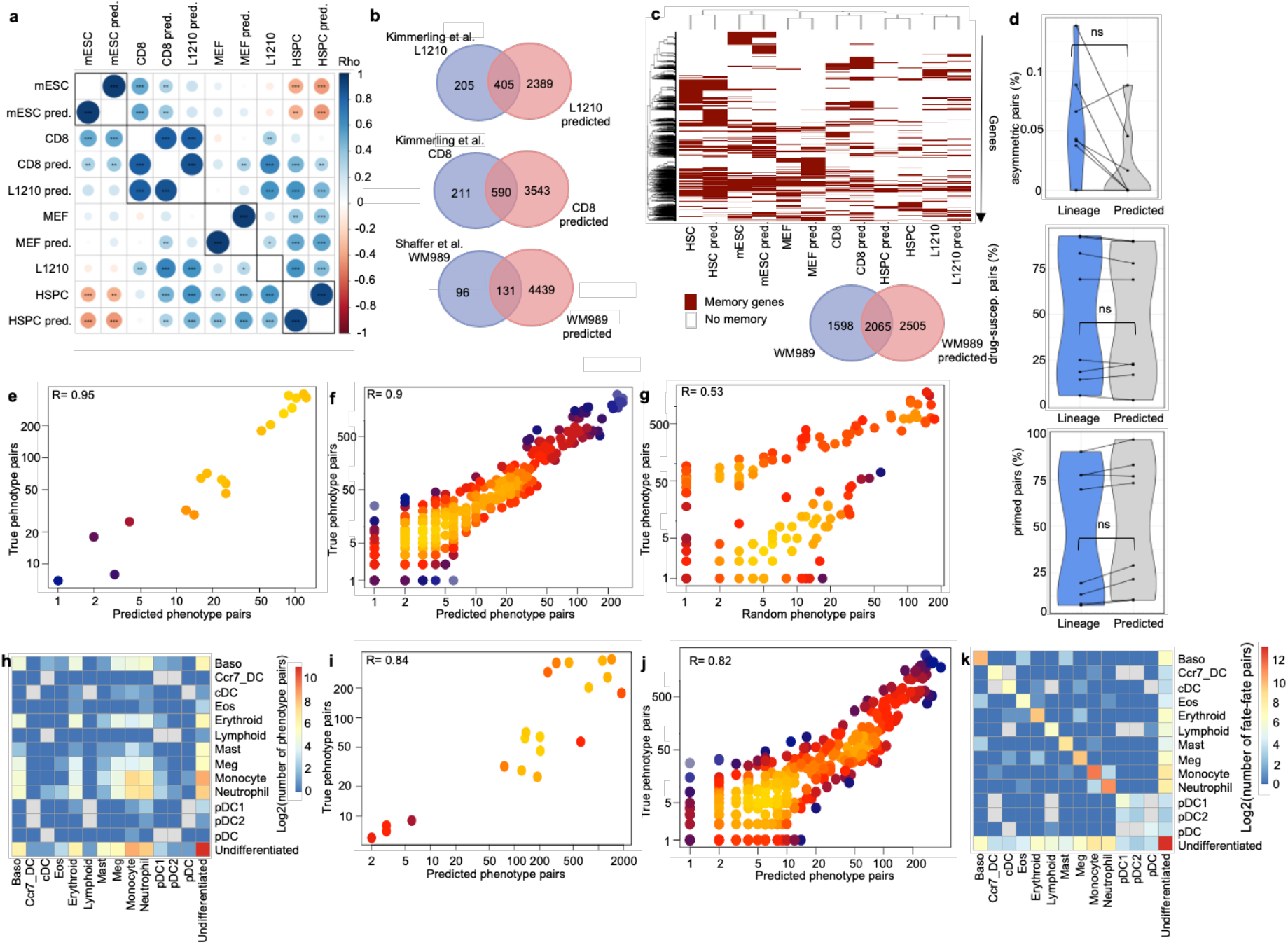
De novo lineage predictions can identify memory genes and cell fate decisions. a) Spearman rank correlation matrix of GO-term enrichment in memory genes called based on true or de novo predicted cell lineages at confidence level 30 in the indicated datasets (Spearman ranks test: !”#$%&’()*+*,-!!”#$%&’()*+*/-!!!”#$%&’()*+**/0+ 1) Venn diagrams showing the overlap of memory genes of CD8, L1210 and WM989 cells from Kimmerling et al. and Shaffer et al. with the memory genes called on de novo predicted lineages at confidence level 50. c) Heatmap (left) showing the overlap of memory genes called on true cell lineages, and memory genes called on de novo predicted lineages as in b across cell types. For the only human cell line analyzed (WM989) overlap is represented in the form of a Venn diagram (right). d) Percentage of asymmetric (left), entirely primed (middle), and entirely drug-susceptible (right) cell pairs in true and de novo lineage predictions (confidence level 50) across all WM989 datasets (n=8). e) Scatterplot showing the number of cell pairs in true and de novo predicted lineages (confidence level 50) in all possible phenotype pair categories across all WM989 datasets as in d. Each dot represents one phenotype pair category in one of the datasets. Coloring represents density in the scatter plot. Spearman rank correlation is given. f) Scatterplot showing the number of cell pairs in true and de novo predicted lineages (confidence level 50) in all phenotype pair categories across all HSPC datasets from 48h and 96h (n=48). g) Scatterplot as in f for random cell pairs. h) Heatmap of the number of cell pairs (log) in random cell lineages in all possible phenotype pair categories in the HSPC datasets as in f. The sum across all datasets is shown. i) Same representation as in e including predicted cell pairs in cells without assigned barcode. j) Same representation as in f including predicted cell pairs in cells without assigned barcode. k) Heatmap of the number of cell pairs (log) in predicted lineages in all possible phenotype pair categories in the HSPC datasets as in j. The sum across all datasets is shown. For d significance was tested using a Mann-Withney U test (p= 0.05).

## Methods

### Cell culture

CGR8 ES cells (Sigma, Cat307032901-1VL) were routinely cultured at 37°C and 5% CO_2_ on dishes coated with 0.1% gelatin type B (Sigma Cat3G9391-100G) in GMEM (Sigma, Cat#G5154-500ML) supplemented with 10% ES cell-qualified fetal bovine serum (Gibco, Cat#16141-079), nonessential amino acids (Gibco, Cat#11140-050), 2 mM l-glutamine (Gibco, Cat#25030-024), sodium pyruvate (Sigma, Cat#S8636-100ML), 100 μM 2- mercaptoethanol (Sigma, Cat#63689-25ML-F), penicillin and streptomycin (Pen/Strep) (BioConcept, Cat#4-01F00-H), homemade leukemia inhibitory factor (LIF), CHIR99021 (Merck, Cat#361559-5MG) at 3 μM and PD184352 (Sigma PZ0181-25MG) at 0.8 μM. Cells were passaged by trypsinization (Sigma, Cat#T4049-100ML) every 2-3 days at a ratio of 1:10. For single cell RNA-seq experiments, cells were switched to N2B27+2i/LIF medium 2 passages beforehand. N2B27+ 2i/LIF medium was composed of a 1:1 mix of DMEM/F12 (Gibco, Cat#11320-033) and Neurobasal (Gibco, Cat#21103-049), supplemented with N2 (Gibco, Cat#17502-001), B27 (Gibco, Cat#17504-001), 1% Pen/Strep (BioConcept, Cat#4- 01F00-H), 2 mM L-glutamine (Gibco, Cat#25030-024), 100 μM 2-mercaptoethanol (Sigma, Cat#63689-25ML-F), LIF, CHIR99021 (Merck, Cat#361559-5MG) at 3 μM and PD184352 (Sigma PZ0181-25MG) at 0.8 μM. Cells were split every 2-3 days at a ratio of 1:10, using accutase (Innovative Cell Technologies 31195) and centrifugation. HEK293T cells were routinely cultured at 37°C and 5% CO_2_ on dishes in DMEM high glucose medium (Gibco #41966) supplemented with 10% Fetal bovine serum (Life technologies #10270-106) and 1% Pen/Strep (BioConcept, Cat#4-01F00-H).

### Lentiviral barcoding library production

The LARRY lentiviral barcoding library ^17^ was purchased from Addgene (https://www.addgene.org/pooled-library/camargo-plarry-egfp-barcoding-v1/). The barcodes of the library are composed of 40 base pairs (bp) with 28 random bp separated by 6 fixed bp doublets, and are located in the 3’ untranslated region of *EGFP* expressed from the *EF-1ɑ* promoter. The library was amplified and lentiviral vector was produced according to the associated protocol with small modifications. Briefly, plasmids were introduced into ElectroMAX Stbl4 Competent Cells (Life technologies #11635018) using MicroPulser Electroporator (Bio-Rad #1652100) EcoRI program. Cells were incubated for 1h at 37°C and spread over 20 large (24.5×14.5 cm) Agar+Ampicillin (Amp) plates. After 24h, colonies were harvested through scraping using pre-warmed LB+Amp. The resulting 1.5 L bacterial culture was incubated at 37°C for 2h and a Maxiprep performed using Qiagen MaxiPlus kit (Qiagen #12963) to a resulting 1 mg of plasmid DNA. Lentiviral vector was produced by transforming the produced LARRY plasmid into HEK293T cells using the Trans-IT 293 transfection reagent (Mirus #MIR2700). For each of eighteen 10 cm dishes of HEK293T cells, 1.5 ml Opti-MEM (Gibco #51935), 16 μg plasmid (8 μg LARRY plasmid, 6 μg PAX2, 2 μg MD2G) were mixed with 45 μl TransIT 293 transfection reagent, incubated for 15-30 min at RT, and added to the cells dropwise. Lentiviral vector particles were harvested 48h and 72h after transfection. Medium was collected and filtered through a 0.45 μm PVDF filter (Millipore #SLHV033RS), and centrifuged in a Beckman Optima XL-80K Ultracentrifuge (Beckman #8043-30-1211) at 20.000 rpm for 1h30 at 4°C. The supernatant was removed, and pellets were resuspended with 100 μl of GMEM with serum and all additives as above but without 2i/LIF. The lentiviral vector preparation was incubated on ice for 30 min, aliquoted and stored at -80°C. Titration of the lentiviral barcoding library was performed on CGR8 cells cultured in GMEM+2i/LIF, with a read-out five days after infection.

### Barcode reference library generation

A reference library was made through sequencing of PCR-amplified barcodes from the LARRY plasmid library in triplicates. 500 ng of LARRY plasmids was taken as input for a two-step PCR using Phusion high fidelity DNA polymerase (Thermo Fisher #F-5305) adapted for LARRY from ^49^. The first step amplifies the barcodes and adds Illumina Read1 and Read2 sequences (5’ACACTCTTTCCCTACACG ACGCTCTTCCGATCTTGTGACGTCACAGGTCGACACCAGTCTCATT3′ and 5’GTGACTGGAGTTCAG ACGTGTGCTCTTCCGATCGAGTAACCGTTGCTAGGAGAGACCATA3′). The second step adds the P5 and P7 flow cell attachment sequences and a sample index of 7 bp (P5 5’AATGATA CGGCGACCACCGAGATCTACACTCTTTCCCTACACGACGCTCTTCCGATCT3′ and P7 5’CAAGCAGA AGACGGCATACGAGANNNNNNNGTGACTGGAGTTCAGACGTGCTCTTCCGATC3′). 200 ng of PCR1 product was taken as input for PCR2. (PCR programs: 98°C 2 min, 15 cycles of 98°C 10 sec, 67.2°C (PCR1), 72°C (PCR2) 30 sec, 72°C 30 sec, followed by final elongation 72°C 5 min and 4°C indefinitely). In between PCR1 and PCR2 and after PCR2, PCR purification was performed using the QIAquick PCR purification kit (Qiagen #28106). The mix of the three samples was sequenced on a MiSeq instrument (Illumina) at the Gene Expression Core Facility of EPFL. Sequencing results were first filtered for a perfect match to the plate index pattern using XCALIBR (https://github.com/NKI-GCF/xcalibr). The resulting read files were filtered for a perfect match to the barcode pattern using customized R scripts. The resulting list and the LARRY “barcode_list” available on Addgene (https://www.addgene.org/pooled-library/camargo-plarry-egfp-barcoding-v1/) were merged and used as reference list.

### mESCs scRNA-seq memory experiment run

CGR8 cells were transduced with the lentiviral LARRY barcoding library to obtain roughly 1% of GFP_+_ cells. 72h after infection, live GFP_+_ cells (Extended Data Fig.1; live stain 1:500 propidium iodide solution (BioLegend #421301)) were sorted on an FACSAria III (BD) at the Flow Cytometry Core Facility of EPFL into Cellcarrier 96-Ultra 96-well plates (PerkinElmer #6055308). Plates were coated with recombinant human E-cadherin-Fc chimera (BioLegend #BLG-779904-25ug) to reduce colony formation and thereby limit the potential impact of paracrine and direct cell signaling in regulating gene expression of related cells ^50^. Coating was performed at 10 μg/ml for 1h at 37°C. After washing the plates with PBS, 1200 cells were sorted into two wells and both were collected after 48h for sequencing. Before collection, cells were imaged on an IN Cell analyzer 2200 (GE Healthcare) at the Biomolecular Screening facility of EPFL. ScRNA-seq library preparation was performed on a 10X Genomics Chromium platform of the Gene Expression Core Facility of EPFL using the SingleCell 3’ Reagent Kit v3.1. The sample was sequenced on a Hiseq4000 instrument (Illumina).

### mESC scRNA-seq data processing and cell lineage inference

ScRNA-seq data was analyzed using 10X Genomics Cell Ranger 5.0.1, Seurat, and customized R and Python scripts. Raw sequencing reads were processed using 10X Genomics Cell Ranger 5.0.1 using default parameters and refdata-gex-mm10-2020-A as reference genome, with or without the *include-introns* option. Cell ranger outputs a unique molecular identifier (UMI) corrected read count matrix. Cells with a percentage of mitochondrial reads between 1.75 and 7.5 and with more than 10,000 reads were further analyzed. Data was normalized to 40.000 reads per cell. Lineage barcodes were extracted from the data using the CellTag pipeline ^18^ available for download at (https://github.com/morris-lab/BiddyetalWorkflow) and adapted to the LARRY barcode design. Briefly the CellTag pipeline extracts reads containing a CellTag motif from the processed, filtered, and unmapped reads BAM files produced in intermediate steps of the 10X Genomics Cell Ranger pipelines. To extract LARRY barcodes, the CellTag motif was changed to “([ACTG]{4}TG[ACTG]{4}CA[ACTG]{4}AC[ACTG]{4}GA[ACTG]{4}GT [ACTG]{4} AG[ACTG] {4})” in all scripts. Barcodes with reads from only one UMI, and without perfect match to the reference library were filtered out. A Jaccard score of >0.7 was used to identify cell lineages. No filtering on the number of barcodes expressed per cell was performed. Lineages were called on unfiltered data. For the mESC dataset, 6 lineages had sizes above 5 cells, which is larger than expected based on cell cluster sizes after 48h of culture and the expected loss of cell lineage members in the preparation for scRNA-seq. They were therefore excluded from further analysis.

### Processing of public data

Data from Biddy et al., 2018 was extracted as BAM file from SRA links specified under GSE99915 (https://www.ncbi.nlm.nih.gov/geo/query/acc.cgi?acc=GSE99915). BAM files were converted back to fastq format using 10X Genomics’ bamtofastq-1.3.2. and Cellranger 5.0.1 was run on the resulting data as described above with or without the *include-introns* option. Cell ranger outputs a UMI corrected read count matrix. Cells were filtered on the percentage of mitochondrial reads and number of reads as indicated in Table 2. Data was normalized to 40.000 reads per cell. Cell lineages were assigned using the CellTag pipeline as specified above with the original CellTag motif of “(GGT([ACTG]{8})GAATTC)”(V1), “(GTGATG([ACTG]{8})GAATTC)”(V2) or “(TGTACG([ACTG]{8})GAATTC)”(V3) and the respective whitelists and sample-specific cell barcodes available at (https://github.com/morris-lab/BiddyetalWorkflow). As in the original publication, cells with >20 or <2 CellTags expressed were not considered for lineage assignment. Lineages were called on unfiltered data using a Jaccard score of >0.7. Cell lineages were called on individual datasets for all analysis of a single time point and across all datasets encompassing clones of the same CellTag library for over-time point analyses.

Data from Kimmerling et al., 2016 is GSE74923_L1210_CD8_processed_data.txt from GEO (https://www.ncbi.nlm.nih.gov/geo/query/acc.cgi?acc=GSE74923). Cell lineages were extracted from the GSE74923_series_matrix.txt file. Data from Weinreb et al., 2020 was downloaded from (https://github.com/AllonKleinLab/paper-data/blob/master/Lineage_tracing_on_transcriptional_landscapes_links_state_to_fate_during_differentiation/README.md). Data from Wehling et al, 2022 was directly obtained from the authors in the form of an unnormalized count matrix and metadata file (will be available on GEO GSE167317 asGSE167317_CountMatrix_Seq5.csv, GSE167317_CountMatrix_Seq4.csv, GSE167317_Metadata_Seq5.csv, GSE167317_Metadata_Seq4.csv). Cells with under 100,000 reads were removed and data was normalized to 40,000 reads per cell. Lineage-information was extracted from the metadata file. Data from Harmange et al, 2022 was downloaded from https://drive.google.com/drive/folders/1-C78090Z43w5kGb1ZW8pXgysjha35jlU?usp=sharing (accessed begin July 2022) in the form of 10×1_Filterd_BatchCor_unnorm_sctrans.rds from experiment one. Lineages were assigned using the corresponding script section in the file 10×1_r1_r2_Analysis_unorm_sctrans.Rmd. During filtering, cells with <3% of mitochondrial reads, and cells with <4,000 reads were removed. Data was normalized to 40,000 reads/cell. Data from Bues et al, 2022 was directly obtained from the authors in the form of a Seurat Object including a normalized count matrix with crypt and organoid annotation. Briefly, organoids were generated by sorting single Lgr5_+_ intestinal stem cells from dissociated organoids into a matrigel matrix. After culture for 3, 4, 5 or 6 days, single organoids were hand-picked and dissociated individually before loading for scRNA-seq. Organoids are derived from 3 batches. Lgr5_+_ intestinal stem cells seeded can be derived from the same organoid. Crypts were collected over 5 batches of 3 pooled mice from 10 mm sections of the ileum. Crypts within the same batch can be derived from the same mouse, but were several mm apart in the ileum.

### General scRNA-seq data representation and analysis

For all scRNA-seq datasets, embeddings were generated using a custom function based on the Pagoda2 R package on exonic, or both intronic and exonic reads. PCA was calculated on the 3000 top variable expressed genes. Subsequently a tSNE was computed on the PCAs top 10 components. Cell cycle was assigned using the cyclone function within scran ^51^. GO-term enrichment analysis was performed using the *topGO* R package. Memory genes and background (all genes detected) were considered to be binary. The GO category “biological process” was used. The top 20 or 100 enriched GOs from each set were visualized as indicated. Velocity analysis was performed using the velocyto package ^33^. Gene expression correlation within cell lineages or random samples for the whole transcriptome or single genes was calculated as Pearson correlation unless otherwise indicated. For comparisons of gene expression correlation within cell lineages over timepoints, pairs of related cells were considered for which one cell was present at day 0 and the other at another time point as indicated. For every lineage shared across timepoints only one cell pair was considered.

### Memory-gene identification and categorization

We defined memory genes as genes with the following characteristics 1) a high correlation in gene expression within cell lineages, 2) a low variability in gene expression between cells of the same cell lineage, 3) a high variability of the mean gene expression in different cell lineages, and being 4) marker genes of cell lineages identified by various Seurat functions. To find genes meeting characteristics 1-3, we called memory genes by comparing the true cell lineage value to the distribution of values from repeated (20X) random sampling of cells. As a variability measure we used the CV_2_ of lineage means for 2) and the intra-lineage CV_2_ for 3). To find marker genes we used the Seurat FindMarker function using a range of *test*.*use* parameters (*bimod, roc, t, negbinom, poisson, LR, MAST*). We then considered the markers with a high sharing across cell lineages, using thresholds defined independently for each *FindMarker()* run to optimize the correlation in gene expression within cell lineages. To identify memory genes using machine learning, we used common dimension reduction methods, mutual information maximizer (MIM) and ANOVA F-test feature selection (ANOVA). Other feature selection techniques frequently used in the literature were also investigated, but MIM and ANOVA outperformed all the other methods. Both MIM and ANOVA work similarly. First, each gene is attributed a memory score either by computing the mutual information or ANOVA F-test of the gene expression across cells. To determine the ideal number of genes (N) to include in the final set, we predicted cell lineages using the top N genes with the highest memory score. Then, each of the resulting clusters was evaluated and the number of genes N with the best precision was chosen as the optimal number of genes. Throughout the main figures memory genes were called using the CV_2_ of lineage means approach. In Extended Data Fig.3 we compare memory genes called using different criteria and methods. Memory genes were called on cell lineages of 3-5 (mESCs) or 2-5 members (all other cell types, and all cell types for machine learning approach). All predicted cell lineages were used when calling memory genes on these. To categorize memory genes into quantitative and qualitative memory genes, we used the *skewness()* function of the R package *moments*. We defined a quantitative memory gene as a gene with a skewness of under 3, and a qualitative memory gene, as a gene with a skewness equal or above 3.

### Cell lineage prediction algorithm in R and python

We developed an iterative clustering approach to predict cell lineages. At every prediction round, the (memory) gene-set used is randomly subsampled to 75% or 50%. The resulting genes are used to cluster the cells of the dataset iteratively. At the start of this iterative clustering, two clusters are generated using *cutree()* function from the dendextend R package and the *hclust()* hierarchical clustering function of the R package stats on the gene expression correlation distance using the agglomeration method ward.D2. In every subsequent iteration round, the cells of each previously defined cluster are reclustered into two clusters using the same procedure. The iteration is ended for clusters meeting a predetermined size of 2-3 cells, or as indicated. 100 predictions are generated based on an each time newly randomly sampled 75% or 50% of the starting (memory) gene set. The result of the 100 iterative predictions are summed in a cell-by-cell matrix. For example, two cells ending up in the same cluster in 30 of the 100 predictions will receive the value of 30. The number of predictions in which a cell pair is found can then be used as a confidence level for the prediction. Precision (TP/(TP+FP)) and sensitivity (TP/(TP+FN)) can be calculated for each confidence level X, by considering only the predictions found in >=X repetitions and comparing them to the true cell lineage relationships. Thereby precision and sensitivity are calculated on cell pairs (Fig.3d for a descriptive scheme of the algorithm). In addition, we also implemented a Python version of the lineage prediction algorithm using the same repetitive clustering approach. As for the R implementation, a random sample containing 75% or 50% of the initial genes is generated 100 times. From this subsampling, the cells are clustered into cell families using the hierarchical clustering function, *ward()* from the SciPy package. We used the Spearman correlation of the gene expression to generate the distance matrix and the ward2 criterion. To restrict the number of cells in each cluster, the *scipy_cut_tree_blanced* library was used. The final lineage assignment and scoring from the 100 independent clusterings was generated as in the R implementation described above. Predictions for datasets under 5 divisions were scored on 1-5 member cell lineages, which are cell lineages for all but the WM989 and mESC dataset. Only for scoring predictions using the machine learning approach, 2-5 member cell lineages were considered. For datasets above 5 divisions, all lineages, or 1-5 member lineages were scored as indicated.

### Parameter selection for predictions

Several parameters can be chosen during the predictions (see Extended Data Fig.7). These are the sampling value (which fraction of genes is selected for each repetition of the iterative clustering), the number of clusters during iterations (how many subclusters are formed at each iteration), the number of repetitions of the iterative clustering (based on which the confidence level is calculated), the size of clusters at which iteration is stopped, and the size of lineages to be predicted. We tested the influence of each independently on the mESC, MEF, CD8, L1210, HSPC, WM989, and HSC datasets starting from values of 100 repetitions, 75% or 50% sampling, 2 clusters formed at each iteration, stopping clustering at 4 cell clusters, and predicted lineages of 2-3 members. These are the default values implemented in the GEMLI prediction function. A lower sampling value increased precision while decreasing sensitivity (Extended Data Fig.7A-B). Changing the numbers of repetitions did only have a small effect on precision and sensitivity values, with higher repetition numbers increasing precision and decreasing sensitivity of the predictions (Extended Data Fig.7C-D). When the number of cells at which clustering is stopped is higher than the maximal lineage size to be predicted, precision values increase and sensitivity decreases (Extended Data Fig.7E-F). Precision was highest when lineage sizes of 2 members were predicted across all datasets, even when datasets encompassed primarily 4 member lineages.

### Gene selection for GEMLI

We selected genes for GEMLI based on the gene expression mean and variability. The variability of gene expression was defined as mean-corrected CV_2_ calculated in the form of the residual of the CV_2_ to a linear fit of CV_2_ and mean expression. Genes for cell lineage predictions were selected based on the quantiles of the mean and variability of gene expression: the 2% highest expressed genes, the 60% most variable among the 10% highest expressed genes, and the 10% most variable among the 40% highest expressed genes: (mean_quantile >= 98) | (mean_quantile >= 90 & variability_quantile >= 40) | (mean_quantile >= 60 & variability_quantile >= 90)).

### Machine learning model to select genes for lineage predictions

To build a classifier capable of discriminating between memory and non-memory genes, we used the mean gene expression and variability as described in the previous section. Memory genes were defined using the CV_2_ of lineage means as described above. The mESCs and MEF cells datasets of 48 and 72h culture time were used for model training, the remaining datasets were kept for model validation. First, the mean expression outliers in each dataset were removed using the Inter quartile range (IQR) method. All the outliers were considered memory genes and were added in the final gene selection. Then, the data was normalized to a range between 0 and 1. Each dataset was preprocessed individually. Different models were trained, including logistic regression with L2 regularization, support-vector machine (SVM) with the radial basis function (RBF) kernel, and a neural network. They were all trained using their sklearn implementation except for the neural network, which was optimized using the Pytorch framework. To optimize the model, we used a balanced version of the binary cross entropy loss and accuracy to account for the fact that the proportion of memory genes is low in all datasets while most machine learning models assume that the data has approximately the same number of samples of both classes. Thus, to resolve this issue, a bigger weight was given to a misclassification of the minority class, the memory genes. To find the optimal hyperparameters, we used a cross-validation and an exhaustive search of all possible hyperparameters for the logistic regression and SVM. For the neural network, we further splitted the data into a training and validation set since cross-validation was deemed too computationally expensive, and used Optuna, a Python package, to perform a random search to find the optimal hyperparameters. The best performing model was the neural network with a validation-balanced accuracy of 0.7108 as compared to 0.552 and 0.697, respectively, for the logistic regression and SVM. Finally, to further assess the final models, we used them to derive a gene set for lineage predictions. The predictions were evaluated as described in the cell lineage prediction algorithm section.

### Assessing the influence of sequencing depth, cell number and lineage structure on lineage predictions

Downsampling of reads and cells was performed on the mESC, MEF and HSPC datasets. For read downsampling, all reads of a given cell were vectorized and subsequently a fraction (66%, 50%, 33%, 10%) of these reads were sampled. Cells were subsampled separately to 66%, 50%, 33%, 10% of the initial cell numbers. Cells without lineage assignment or being the only member of a cell lineage were subsampled directly to the desired fraction. Cells that were members of cell lineages with several members were subsampled together with their respective cell lineage members. The third parameter whose influence on predictions was analyzed was the lineage structure of datasets. Importantly, lineage assignment using cellular barcodes in scRNA-seq data is not standardized with respect to the thresholds in barcode similarity based on which cells are assigned to the same lineage. Furthermore, some lineage assignment pipelines disregard cells which they cannot link to any other cell, while others record them as lineages with a single member. Single-member lineages can thereby be true results of different experimental setups, or technical artifacts of barcode-based lineage assignments. Lineage structure was changed by either including all cells without barcode assignment as single member lineages in the mESC dataset, or excluding single member lineages from the HSPC dataset before GEMLI predictions and scoring.

### Cell fate analysis

To analyze how well GEMLI can recapitulate cell fate decisions, we analyzed HSPC data from Weinreb et al. and WM989 data from Harmange et al.. In Weinreb et al., HSPCs are cultured in myeloid differentiation conditions and are annotated according to their phenotype as basophil, eosinophil, erythroid, lymphoid, megakaryocyte, monocyte, neutrophil, Mast cell, DC (pDC, cDC, pDC1, ccr7_DC) or undifferentiated cells. Only certain phenotype combinations of related cells exist and all have different prevalence, meaning that HSPCs are restricted in their fate decisions. For all cell pairs of true and GEMLI-predicted cell lineages, the annotated phenotypes from Weinreb et al. were extracted from the published metadata. For each possible phenotype pair category (for example erythroid-eosinophil or undifferentiated-Mast cell) the number of cell pairs in true and GEMLI predictions was determined and summed for all datasets. As 48h datasets encompassed mainly undifferentiated cells, all 96h HSPC datasets were also analyzed (total of 48 datasets; Supplementary table 1). In Harmange et al., each WM989 cell is annotated as drug-susceptible or primed (drug-resistant). Members of each cell lineage are either all in one of these two states or distributed between these two states (asymmetric cell pairs), with the latter implying that cells recently switched fate. We extracted the annotation for each cell from the published Seurat object and compared the number of true and de novo predicted related cell pairs being entirely drug-susceptible, primed or distributed between the two states. DEG for asymmetric cell pairs and entirely drug-susceptible or primed cell pairs were called as in the original publication, using the *FindMarkers()* function of Seurat with parameters min.pct=0.05 and logfc.threshold=0.1.

## Data availability

Data will be made available on GEO. The R package for lineage predictions and memory gene identification and its documentation are available on Github at https://github.com/UPSUTER/GEMLI.

## Acknowledgments

We would like to acknowledge the following funding sources: Swiss National Science Foundation grant#CRSK-3_195097, and Novartis Foundation for Medical-Biological Research grant#20A023 to DMS., OE och Edla Johanssons Research Foundation to MT and ASE, The Swedish Research Council (VR 2021-06112 and 2020-01480), Knut and Alice Wallenberg Foundation (KAW 2016.0123), Ragnar Söderberg Foundation (S12/20) and Karolinska Institutet to VP and MT. We also would like to thank the Gene Expression Core Facility, Flow Cytometry Core Facility, Biomolecular Screening facility and Scientific IT & Application Support of EPFL. Computational analysis was partially performed on resources provided by the SNIC through UPPMAX partially funded by the Swedish Research Council (2018-05973). We thank Theo Maffei and Imane Ben M’Rad as students of the EPFL Machine learning Master course CS-433, as well as Bart Deplancke, Gioele La Manno, Nicholas Phillips, Vincent Gardeux, Johannes Bues, Jörn Pezoldt, Nikolay Oskolkov, Gustavo Jeuken, and the Friedländer, Kutter and Pelechano labs for their helpful input.

## Contributions

A.S.E. and D.M.S conceived and designed the project. D.M.S and V.P supervised the project. A.S.E. collected the ES cell data. A.S.E and M.T. analyzed the data and developed GEMLI. M.T. wrote the GEMLI R package. A.A.D. performed the machine learning analysis and implemented a GEMLI python version. A.S.E. and D.M.S wrote the manuscript with contributions from all authors.

